# EWSR1 affects PRDM9-dependent histone 3 methylation and provides a link between recombination hotspots and the chromosome axis

**DOI:** 10.1101/325282

**Authors:** Hui Tian, Timothy Billings, Michael Walker, Pavlina M. Petkova, Christopher L. Baker, Petko M. Petkov

## Abstract

Meiotic recombination in most mammals requires recombination hotspot activation through the action of the histone 3 lysine-4 and lysine-36 methyltransferase PRDM9 to ensure successful double-strand break initiation and repair. Here we show that EWSR1, a protein whose role in meiosis was not previously clarified in detail, binds to both PRDM9 and pREC8, a phosphorylated meiosis-specific cohesin, in male meiotic cells. We created a *Ewsr1* conditional knockout mouse models to deplete EWSR1 before the onset of meiosis, and found that absence of EWSR1 causes meiotic arrest with decreased histone trimethylation at meiotic hotspots, impaired DNA double-strand break repair, and reduced crossover number. Our results demonstrate that EWSR1 is essential for promoting PRDM9-dependent histone methylation and normal meiotic progress, possibly by facilitating the linking between PRDM9-bound hotspots and the nascent chromosome axis.

**Author Summary:** In most mammals, including humans and mice, genetic recombination initiates when the meiosis-specific protein PRDM9 binds specific DNA sequences, known as hotspots, at the beginning of the extended prophase I of meiosis, and activates them by trimethylating histone 3 at lysine-4 and lysine-36 on nearby nucleosomes. Although this activation of hotspots is believed to occur on the chromatin loops, the subsequent double-strand break formation and repair occur on a proteinaceous structure known as the chromosome axis. We now show that Ewing sarcoma RNA binding protein 1 (EWSR1) is a key player in early recombination events, binding to PRDM9, promoting PRDM9-dependent histone methylation, and facilitating the linking between PRDM9-bound hotspots and the nascent chromosome axis through the meiosis-specific cohesion REC8. As a result of these activities, EWSR1 assures sufficient numbers of properly positioned crossovers in each meiosis.

## Introduction

Correct genetic recombination during meiosis is required for the production of fertile, euploid gametes and the creation of new combinations of parental alleles [1–3]. In most mammals, including humans and mice, recombination is enriched at defined genomic sites, termed hotspots, whose positions are determined by the meiosis-specific protein PR/SET Domain 9 (PRDM9) [4–6]. Meiotic recombination initiates at leptonema, the first stage of the prolonged prophase I of meiosis, when PRDM9 binds to hotspot sequences with its zinc finger domain and trimethylates histone 3 at lysine 4 (H3K4me3) and lysine 36 (H3K36me3) with its PR/SET domain [4–8]. Double-stranded DNA breaks (DSBs) then form at some of the activated sites by the action of the topoisomerase-like protein SPO11 [3, 9]. The strand cleaved by SPO11 is then resected [3] and the resulting single-strand ends are used to establish contact between homologous chromosomes [10]. The resulting recombination intermediates eventually resolve to form crossovers or non-crossovers, with a strong preference for non-crossovers [11–13]. The number and positioning of the PRDM9-trimethylated sites, DSBs, and crossovers are under stringent control. In C57BL/6J (B6) male mice, it is estimated that in an average meiosis ∼4,700 hotspots are modified by PRDM9 [14], but of these, only ∼200-300 are used for DSB formation [15, 16], and in turn these are repaired to produce mostly non-crossovers and only 22-24 crossovers [17]. This strict regulation of meiotic recombination events at each step is required to ensure accurate progression of meiosis [2, 16]. The determination of whether or not a PRDM9 binding site is used for recombination initiation is affected by multiple factors, including allelic competition [18, 19] and local chromatin structure [20].

The complexity of the process of meiosis, and the stringent controls in place, require the involvement of multiple proteins. It is believed that PRDM9 binds to DNA in the chromatin loops. However, many of the proteins involved in DSB formation and repair that are currently known [9, 11, 21–24], are physically located on the chromosome axis, a proteinaceous structure which begins to form at the outset of meiosis, when cohesin protein complexes establish the core of the chromosome axis [25], and subsequently assemble the synaptonemal complex (SC) linking the two homologous chromatids together in meiotic prophase I.. We have little understanding of how the recombination machinery recognizes PRDM9-activated hotspots, what decides which of these will be chosen for DSB formation, and how DSBs are associated with the chromosome axis elements [26]. We recently identified Ewing sarcoma RNA binding protein 1 (EWSR1) as a binding partner of PRDM9 during meiotic prophase I through its C-terminal domain, and showed that PRDM9 also interacts with elements of the chromosome axis including pREC8, a phosphorylated meiotic cohesin protein, and the synaptonemal complex proteins SYCP3 and SYCP1 [27].

In wild-type mice, EWSR1 is expressed in most tissues and cell types, including both Sertoli cells and germ cells in seminiferous tubules [27]. Systemic knockout of *Ewsr1* in mice results in high postnatal lethality, defects in pre-B cell development, premature cellular senescence, hypersensitivity to ionizing radiation, and sterility in any surviving mice [28]. Therefore, to further evaluate the function of EWSR1 specifically in meiotic recombination *in vivo*, we created a conditional knockout mouse model in which *Ewsr1* is deleted in germ cells at the onset of meiosis. Here, in experiments using this model we show that EWSR1 plays important roles in early meiotic prophase, promoting PRDM9-dependent histone methylation at hotspots and providing a physical link between activated hotspots and pREC8-containing meiosis-specific cohesin complexes. These actions of EWSR1 further affect the subsequent hotspot choices for DSB formation and crossover resolution.

## Results

### EWSR1 links PRDM9 with the chromosome axis elements through pREC8-containing cohesin complexes

Based on our previous finding showing that PRDM9 binds directly to EWSR1 and strongly but indirectly to pREC8, the meiosis-specific kleisin cohesin subunit [27], we used co-immunoprecipitation (co-IP) to test whether EWSR1 mediates the connection between PRDM9 and pREC8-containing chromosome axis cohesins. First, we detected a strong co-IP signal between EWSR1 and pREC8 (Fig. 1A). This interaction was retained after DNase I treatment, suggesting that the interaction between EWSR1 and pREC8 does not depend on their association with DNA. In contrast, the second meiosis-specific kleisin subunit, RAD21L, showed very weak signal in EWSR1 co-IP, which became almost undetectable after DNase I treatment (Fig. 1A).The SC proteins SYCP3 and SYCP1 (Fig. S1 and [27]) did not show any interaction with EWSR1.

**Fig 1.**
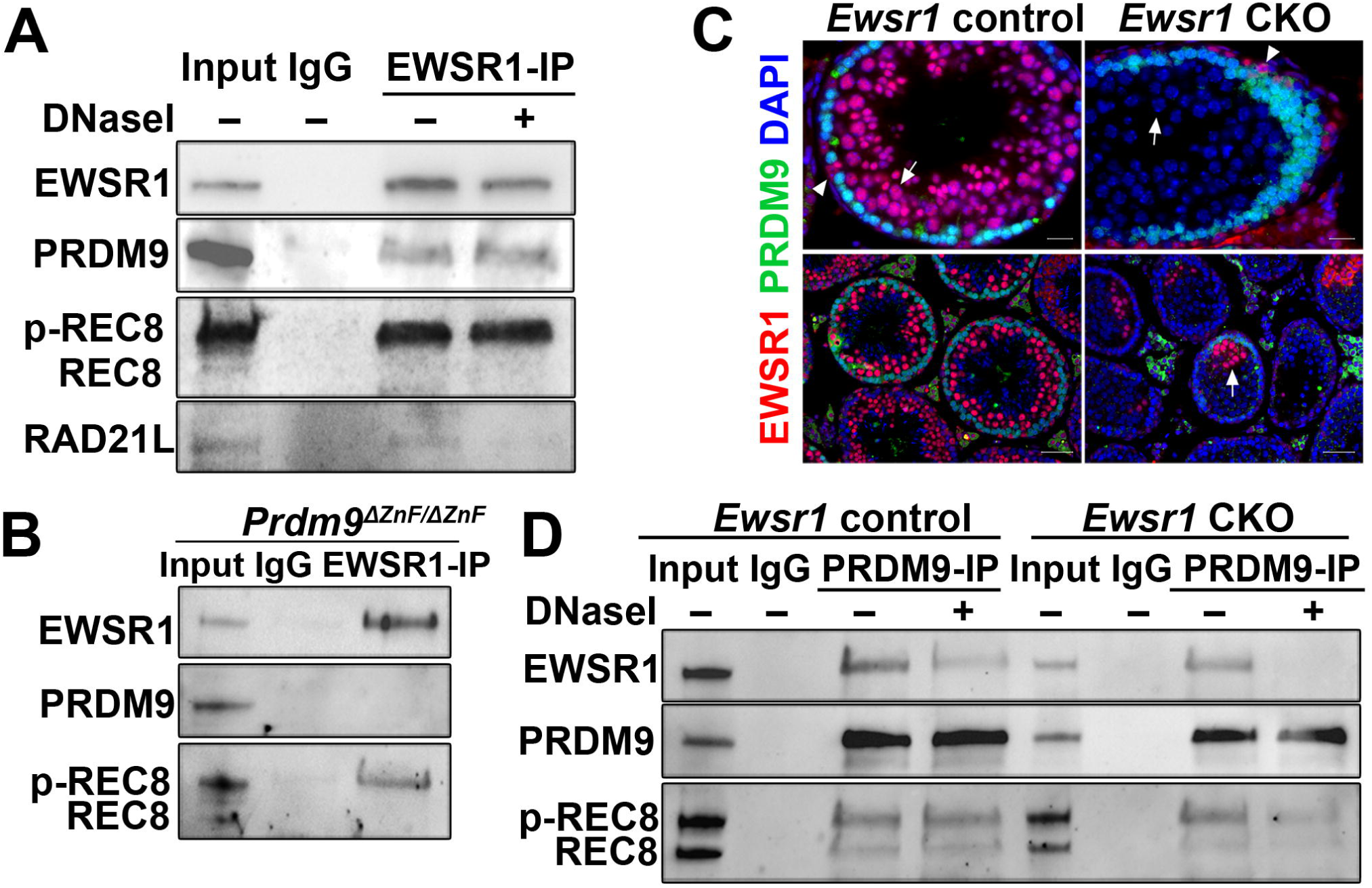
EWSR1 is co-expressed with PRDM9 and interacts with it and with meiotic cohesin protein pREC8 in spermatocytes. (A) Co-IP with anti-EWSR1 in 14-dpp control testis extract stained for PRDM9 and cohesin proteins from 14 dpp B6 spermatocytes. In each blot, lane 1 – input; lane 2 – co-IP with non-immune IgG; lane 3 – co-IP with anti-EWSR1 untreated with DNAse I; lane 4 – co-IP with anti-EWSR1 treated with DNAse I. (B) Co-IP with anti-EWSR1 and anti-PRDM9 in 14 dpp *Prdm9*^ΔZnF/ΔZnF^. Lane 1 – input; lane 2 – co-IP with non-immune IgG; lane 3 – co-IP with anti-EWSR1. (C) EWSR1 (red) and PRDM9 (green) staining of *Ewsr1* control and CKO seminiferous tubule sections. Arrows indicate spermatocytes; arrowheads indicate Sertoli cells; arrow in lower panels indicates EWSR1 is leaky expressed in pachytene cells in CKO. Scale bar in upper panels, 20 µm; scale bar in lower panels, 50 µm. (D) Co-IP with anti-PRDM9 in 14 dpp *Ewsr1* control (lanes 1-4) and CKO (lanes 5-8) testis extracts. In each blot, lanes 1 and 5 – input; lanes 2 and 6 – co-IP with non-immune IgG; lanes 3 and 7 – co-IP with anti-EWSR1 untreated with DNAse I; lanes 4 and 8 – co-IP with anti-EWSR1 treated with DNAse I.

In *Prdm9* knockout (*Prdm9^tm1Ymat^* [29], referred to as *Prdm9^-/-^*) spermatocytes, the interaction between EWSR1 and pREC8 is preserved. However, in *Prdm9^tm3.1Kpgn^* mutant mice, which express a PRDM9 fragment that lacks its zinc finger domain and is thus unable to bind DNA or trimethylate histone 3 at hotspots (hereafter *Prdm9^ΔznF/ΔznF^*), PRDM9 shows no interaction with pREC8, and shows dramatically reduced interaction with EWSR1 even though its EWSR1-binding N-terminal domain is preserved [27]. We hypothesized that the DNA binding activity of PRDM9 is required to retain the association between EWSR1, PRDM9, and pREC8. To test this hypothesis, we performed co-IPs with EWSR1 in spermatocytes of *Prdm9*^ΔznF/ΔznF^ mice. In this mutant, EWSR1 co-immunoprecipitated pREC8 but not PRDM9 (Fig. 1B and [27]). These results suggest that the interaction between EWSR1 and pREC8 is independent of the presence of PRDM9 or its binding to DNA. However, the results also show that loss of PRDM9’s DNA binding activity disrupts the interaction between PRDM9 and EWSR1, resulting in disassociation between PRDM9 and the cohesin complexes.

To further investigate the interactions between EWSR1, PRDM9 and pREC8, we created *Ewsr1* conditional knockout mice (*Ewsr1^loxp/Δ;Stra8-iCre^*, hereafter referred to as *Ewsr1* CKO) by deleting *Ewsr1* from spermatocytes at the onset of meiosis (before pre-leptonema) using *Stra8-iCre*. As expected for successful deletion, the EWSR1 signal was absent in spermatocytes only in the *Ewsr1* CKO testis (Fig. 1C, arrow in upper panels). Spermatogonia and Sertoli cells were not affected (Fig. 1C, arrowhead in upper panels. A small percentage of spermatocytes (<10%) showed incomplete EWSR1 excision (Fig. 1C, arrow in lower right panel). We then compared the interactions involving the three proteins in the CKO and control testes by performing co-IP with PRDM9. In the heterozygous control spermatocytes (*Ewsr1^loxp/+;Stra8-iCre^*, hereafter referred to as *Ewsr1* control), PRDM9 bound to both EWSR1 and pREC8 with or without DNase I treatment (Fig. 1D, left four lanes). In CKO spermatocytes, PRDM9 showed some interaction with pREC8 without DNase I treatment, but this was dramatically reduced after DNase I treatment (Fig. 1D, lower panel, right four lanes). In CKO spermatocytes, PRDM9 interacted with EWSR1 without DNase I treatment (Fig. 1D, upper panel, lane 7) but did not interact with EWSR1 after DNAse I treatment (Fig. 1D, upper panel, lane 8). The latter result is expected, as EWSR1 is not expressed in these CKO spermatocytes. The interaction between the two proteins detected in the absence of DNAse I treatment Fig. most probably stems either from the association between EWSR1 from somatic cells and DNA after cell lysis, or from incomplete EWSR1 excision in the small percentage of spermatocytes (Fig. 1C).

Together, these data indicate that EWSR1 could mediate the association between PRDM9 and the chromosome axis cohesin complexes via binding to pREC8.

### *Ewsr1* conditional knockout (CKO) mice are sterile with meiotic arrest

We further investigated the requirement of EWSR1 for proper meiosis progression and fertility. First, we determined the fertility parameters of *Ewsr1* CKO mice. Similarly to the systemic knockout mice [28], *Ewsr1* CKO mice were sterile, while control matings produced a normal number of viable pups (Fig. 2A). Loss of *Ewsr1* in spermatocytes led to a significant reduction in testis size compared to controls in adult mice; the testis weight index was markedly lower in CKO adult mice than in control adult mice (Fig. 2B), but not in juvenile mice at 14 or 18 days postpartum (dpp) (Fig. S2A). Histologically, *Ewsr1* deficiency resulted in loss of spermatocytes in seminiferous tubules beyond prophase I and loss of post-meiotic germ cells (Fig. 2C, upper panels). Consequently, no mature sperm were observed in the CKO epididymis (Fig. 2C, lower panels). There was also a marked increase in the number of apoptotic germ cells in CKO compared to control testes (Fig. 2D and E).

**Fig. 2.**
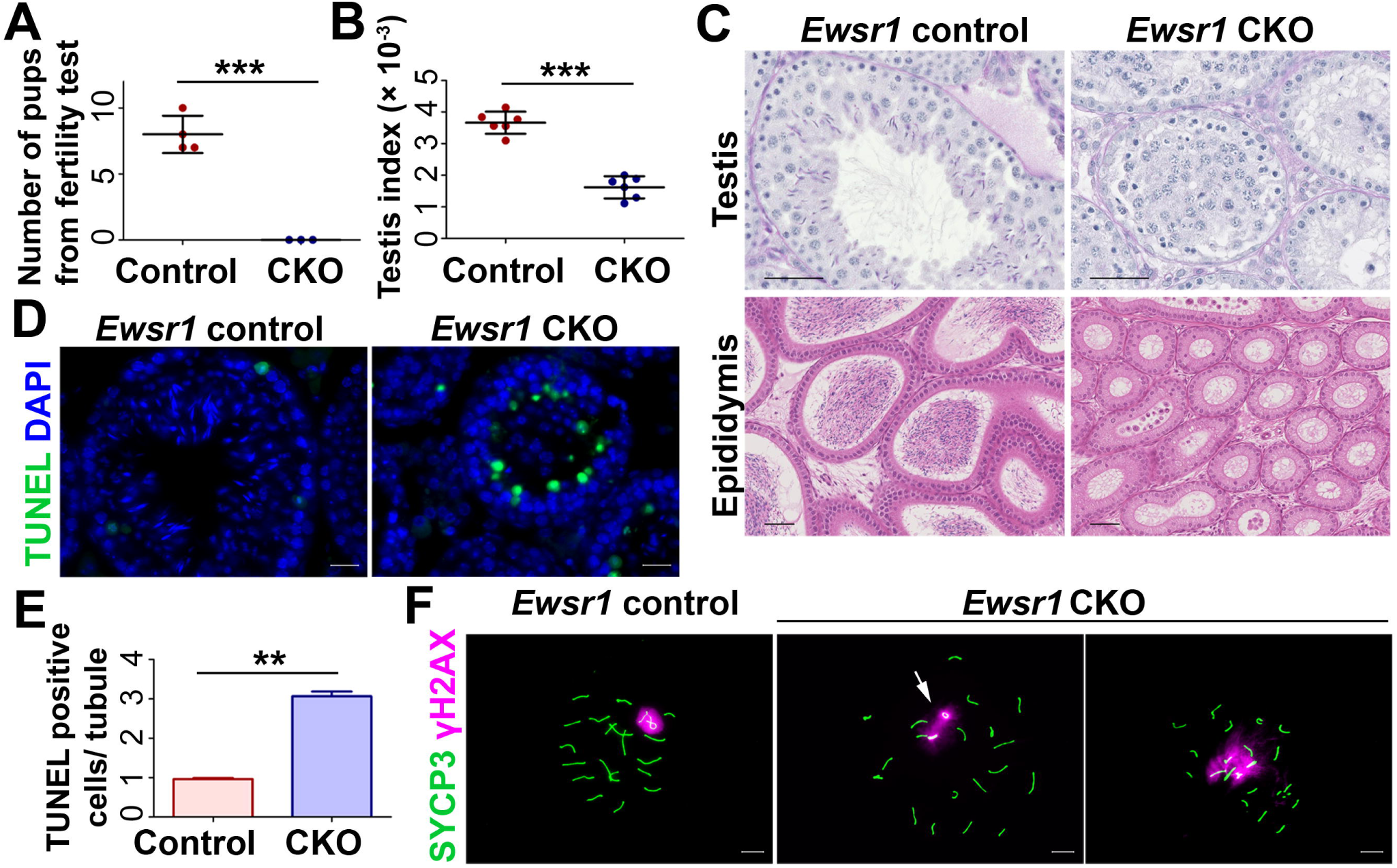
*Ewsr1* CKO mice are sterile with meiotic arrest and chromosomal asynapsis in spermatocytes. (A) Fertility test in *Ewsr1* control (n = 4) and CKO (n = 3) male mice. (B) Testis index (testis weight/body weight) in *Ewsr1* control and CKO mice. (C) Top panels - PAS staining of seminiferous tubule sections. Bottom panels - H&E staining of epididymis sections. Left panels - control; right panels - *Ewsr1* CKO (right). Scale bar, 50 µm. (D and E) Apoptosis in the *Ewsr1* CKO and control testes. (D) TUNEL staining in *Ewsr1* control and CKO testis. Scale bar, 50 µm. (E) Apoptotic cell number quantitated as TUNEL positive cell number per tubule cross section in *Ewsr1* control and CKO. (F) Spermatocyte chromosome spreads of control and CKO stained for γ (magenta) and SYCP3 (green). Arrow in middle panel shows XY chromosomes asynapsis. Arrow in right panel shows autosome and XY asynapsis. Scale bar, 10 µm. Bars in (A), (B) and (E) represent mean ± SD. ** *p* < 0.01, *** *p* < 0.001, by Student’s *t*-test.

We then examined meiotic progression in CKO testis, using spermatocyte spreads. Meiosis appeared to be blocked at pachynema/diplonema in CKO mice. Although the proportions of late pachytene spermatocytes, marked by H1t [30]Fig., were reduced in CKO compared to control testes (Fig. S2B, *p* < 0.05), the proportions of leptotene and zygotene spermatocytes were not significantly different in CKO and control testes (*p* = 0.21, *Chi-*square, Fig. S2C).

Next, we examined the formation and repair of DSBs using phosphorylated H2AX (γH2AX) as a marker of unrepaired DNA lesions. In normal meiosis, the γH2AX signal is seen throughout the nucleus in leptotene spermatocytes when DSBs are occurring. The repair of DSBs results in loss of γH2AX signal in the autosomes by pachynema, when the signal becomes restricted to the sex body [31]. In CKO spermatocytes, the γH2AX signal was detected normally during leptonema and zygonema, and at pachynema most of the signal was restricted to the sex body as in wild type controls (Fig. S2D). However, ∼24.3% of CKO sex chromosomes were not attached to each other, compared to 0.7% in *Ewsr1* control spermatocytes (*p* < 0.05, Fig. 2F, middle panel, Fig. S2D, arrow) In addition, ∼8.5% of CKO pachytene spermatocytes showed remaining γH2AX signal on autosomes, compared to ∼0.9 % in controls (*p* < 0.05). These results suggest asynapsis at autosomes (Fig. 2F, right panel). To further examine the effect of EWSR1 loss on sex body formation and synapsis, we used ATR, BRCA1 and HORMAD1 as markers of both sex bodies and unsynapsed chromosome axes. Round–spread ATR signal covering the sex chromosomes and BRCA1 signal along the XY axes were observed on the sex bodies of both CKO and control pachytene spermatocytes (Fig. S2E), but, as with γH2AX staining, larger numbers of unsynapsed XY chromosomes were observed in CKO than in control spermatocytes (Fig. S2E, arrows). HORMAD1 staining also indicated XY chromosomes asynapsis in CKO spermatocytes, with no such asynapsis in control spermatocytes. There was also a marked increase in autosomal asynapsis in CKO compared to control spermatocytes (Fig. S2E, arrow).

Together, these results suggest that depletion of EWSR1 in spermatocytes affects normal DSB repair and chromosome synapsis.

### PRDM9 and its interactor proteins are expressed normally in the absence of EWSR1

To investigate whether the meiotic defects caused by *Ewsr1* germ cell deletion are PRDM9-related, we first determined whether the expression and/or localization of PRDM9 and its interactors are affected in the CKO spermatocytes. Normally, PRDM9 expression is restricted to the nucleus at leptonema and zygonema [32]. In the absence of EWSR1, there was no change in PRDM9 expression pattern (Fig. S3A, upper two rows) or its protein level (Fig. 1C) in 14 dpp CKO spermatocytes. We examined the other PRDM9 interactors we identified previously [27], and, similarly, found no difference in protein levels of EHMT2 and the chromosome axis proteins REC8 and SYCP3, in CKO testes compared to controls (Fig. S3A, third-fifth rows), and the pattern of EHMT2 or CDYL localization was the same in spermatogonia and spermatocytes of CKO and controls (Fig. S3B).

Together, these data indicate that EWSR1 is not involved in regulation of the expression of PRDM9 or its interactors, or their nuclear localization during spermatogenesis.

### H3K4 and H3K36 trimethylation levels at hotspots are reduced in the absence of EWSR1

Next, we investigated whether EWSR1 affects hotspot activation by PRDM9 by comparing the patterns of H3K4me3 and H3K36me3 marks in control, *Prdm9^-/-^*, and *Ewsr1* CKO testes. Immunostaining of spreads of wild type B6 mice showed abundant H3K4me3 signal present at leptonema and zygonema but very little at pachynema (Fig. 3A, upper panel, first row, and [33]), a pattern that matches the expression of PRDM9 (Fig. 3A, upper panel, first row, and [27]). In contrast, H3K4me3 signal was low in both leptonema/zygonema and pachytene-like cells in *Prdm9^-/-^* spermatocytes (Fig. S3C, second row), indicating that the robust H3K4me3 signal in wild type leptonema and zygonema is mostly PRDM9-dependent. In *Ewsr1* CKO spermatocytes, H3K4me3 signal at leptonema and zygonema was reduced when compared to controls (Fig. 3A lower panel, third row), in a manner similar to those in *Prdm9^-/-^* spermatocytes.

**Fig 3.**
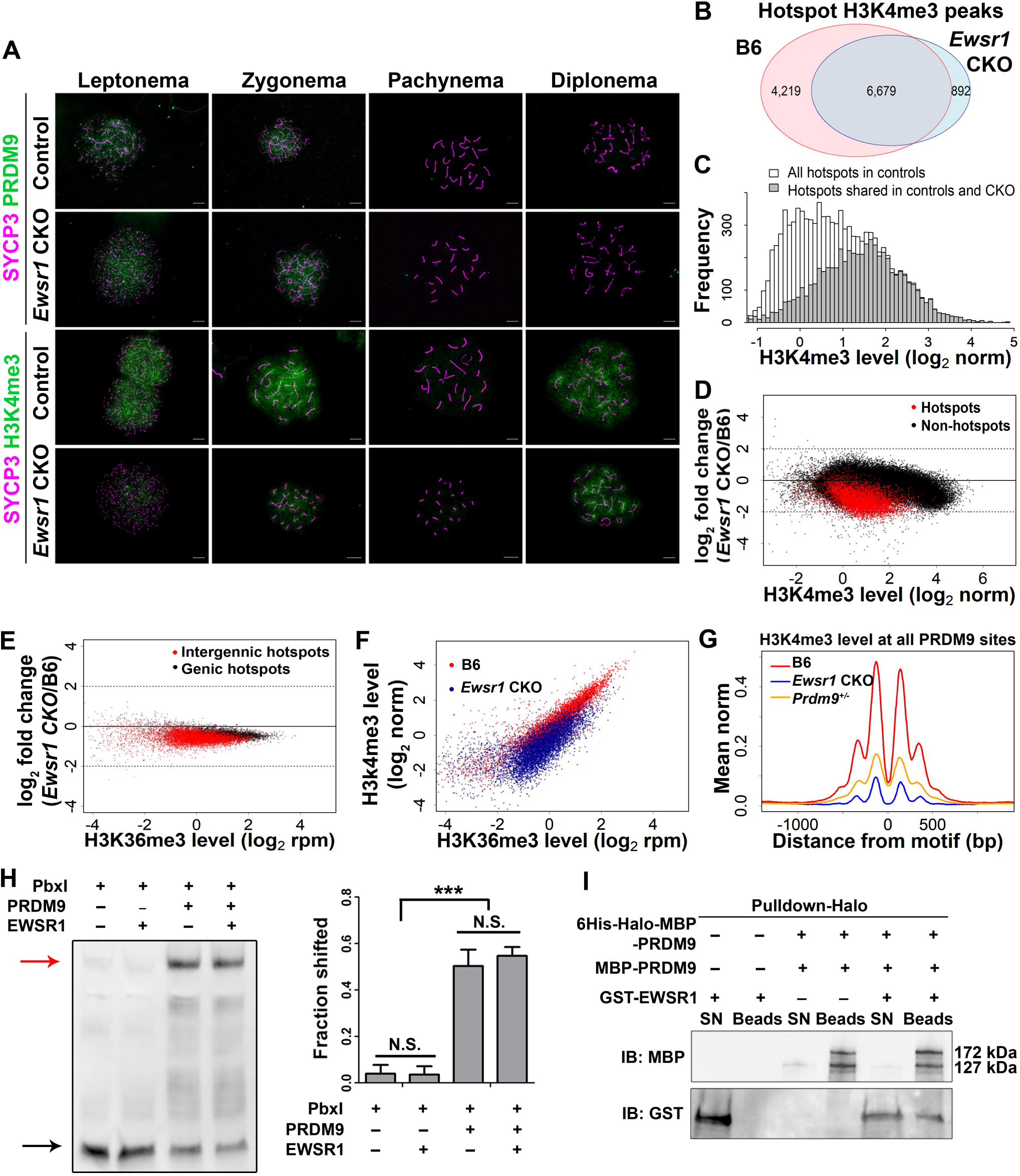
H3K4me3 and H3K36me3 activities at hotspots are reduced in *Ewsr1* CKO. (A) Chromosome spreads on *Ewsr1* control and CKO stained for PRDM9 (green), H3K4me3 (green) and SYCP3 (red). Scale bar, 10 µm. (B) Venn diagram of H3K4me3 ChIP-seq hotspot peaks in control (pink) and CKO (blue) spermatocytes. (C) Distribution of H3K4me3 hotspot frequency levels in B6 control (white) and *Ewsr1* CKO (grey). (D) MA-plot of shared H3K4me3 peaks in B6 and *Ewsr1* CKO mice. Red dots, PRDM9-defined H3K4me3 hotspots, n = 6,679; black dots, H3K4me3 peaks at non-hotspots, n = 54,434. (E) MA-plot of H3K36me3 reads in PRDM9-defined H3K4me3 hotspots from *Ewsr1* CKO and control spermatocytes. Red dots, hotspot in intergenic region, n = 6,082. Black dots, hotspot in genic region, n = 4,791. (F) Correlation between H3K4me3 and H3K36me3 level at hotspots in B6 control (red dots) and CKO mice (blue dots). (G) Aggregation plots of H3K4me3 signal in B6 (red lines), *Prdm9^+/-^* (yellow lines) and *Ewsr1* CKO (blue lines) spermatocytes, centered at the PRDM9 binding site. Left, mean rpm counts. (H) EMSA with MBP-PRDM9 and Pbx1 hotspot in the presence or absence of 6His-HALO-EWSR1. Red arrow, shifted band; black arrow, unbound fragment. Right, quantification of shifted fraction. Data represent mean ± SD. *** *p* < 0.001, N.S., not significant, by two-way ANOVA. (I) Pull-down assay with purified 6HisHALO-MBP-PRDM9, MBP-PRDM9 and GST-EWSR1. SN, supernatant. Beads, elution from beads.

Strong H3K36me3 signal was detected in leptonema, zygonema, and diplonema in control mice, reflecting its presence both at hotspots and at actively transcribed gene bodies (Fig. S3D, upper panel). In *Ewsr1* CKO spermatocytes, H3K36me3 signal was reduced when compared to controls in leptonema and zygonema, but not in diplonema (Fig. S3D, lower panel).

To test whether loss of EWSR1 affects the methyltransferase activity of PRDM9 at hotspots, we performed H3K4me3 ChIP-seq on 14-dpp control and CKO testes. We did each of these experiments in two replicates, which showed excellent correlation (*r*=0.99, Fig. S3E). We found 10,898 PRDM9-dependent H3K4me3 peaks in controls, and only 7,571 peaks in CKO spermatocytes (FDR < 0.01, Fig. 3B) using the detection criteria described previously [20]. Over 88.2% of the hotspot peaks in CKO spermatocytes were present in the controls (6,679 out of 7,571). Conversely, about 38.7% of the hotspot peaks in control spermatocytes (4,219 out of 10,898) were missing in CKO spermatocytes (Fig. 3B). The missing hotspots were those that had relatively low H3K4 trimethylation levels in controls (Fig. 3C), and for the vast majority of shared hotspots, H3K4me3 levels were generally lower in CKO spermatocytes than at the corresponding location in control spermatocytes (Fig. 3D, red points. *p* < 2×10^-16^, Wilcoxon rank sum test, and Fig. S3F). In contrast, H3K4 trimethylation at non-hotspots was not significantly different in CKO versus control spermatocytes (Fig. 3D, black points). The reduction in the H3K4 trimethylation levels at hotspots following loss of *Ewsr1* in spermatocytes was not due to depletion of leptotene/zygotene spermatocytes in CKO testes; in fact, the proportion of zygotene spermatocytes in CKO testes was slightly increased following loss of *Ewsr1* (Fig. S2C).

PRDM9-dependent H3K36 trimethylation at hotspots [7, 34], also showed reduced levels in CKO compared to control spermatocytes (Fig. 3E. *p* < 2×10^-16^, Wilcoxon rank sum test). This reduction of H3K36 trimethylation levels was stronger in hotspots in intergenic regions than those in actively transcribed genes (Fig. 3E, *p* < 2×10^-16^, Wilcoxon rank sum test), Fig.Fig.likely due to the additional presence of H3K36me3 background associated with transcription. The reductions in H3K4me3 and H3K36me3 levels at all hotspots in *Ewsr1* CKO spermatocytes correlated with each other (Fig. 3F).

Together, these results indicate that EWSR1 affects PRDM9-dependent H3K4 and H3K36 trimethylation in spermatocytes.

### EWSR1 does not affect PRDM9-hotspot DNA binding or PRDM9 multimerization *in vitro*

We next investigated whether loss of *Ewsr1* affects the overall levels of PRDM9-dependent H3K4 trimethylation. The mean H3K4me3 level at all PRDM9-dependent sites was much lower in CKO spermatocytes Fig.than in controls Fig. or *Prdm9* heterozygous (*Prdm9^+/-^*) spermatocytes [35] (Fig. 3G).

Two possible explanations of the reduced H3K4me3 level at hotspots in CKO spermatocytes are that EWSR1, which strongly binds PRDM9 *in vitro* [27], could affect PRDM9 binding to hotspot DNA or its multimerization [34–36]. We tested the effect of EWSR1 on PRDM9 binding to hotspot DNA by electrophoretic mobility shift assay (EMSA) using two hotspot DNA fragments (Pbx1 [37] and Chr16-66.9Mb) and *E.coli*-expressed purified EWSR1 and PRDM9. For each of the two hotspot fragments, binding of PRDM9 to hotspot DNA was unaffected by status (presence or absence) of EWSR1 (Fig. 3H and Fig. S3G). Next, we investigated whether EWSR1 is necessary for PRDM9 multimerization. We first tested directly whether full-length PRDM9 molecules can form multimers *in vitro*, using PRDM9 fused to two different tags, 6HisHALO-MBP and MBP alone. The two fused proteins differ in their molecular weight (172 kDa and 127 kDa, respectively). PRDM9 molecules did bind one another (Fig. S3H), and this interaction was not mediated by the MBP tag (Fig. S3I). We then examined the impact of EWSR1 on this interaction and showed that EWSR1 strongly interacted with PRDM9 (Fig. 3I, lower panel, lane 6, and [27]), but did not affect the interaction between the PRDM9 molecules *in vitro* (Fig. 3I, upper panel, lanes 4 and 6). These results show that PRDM9 can form multimer complexes independently of its interaction with EWSR1, at least *in vitro*.

In summary, loss of EWSR1 affects the efficiency of H3K4/K36 trimethylation at meiotic hotspots *in vivo*, markedly decreasing their number and activity, but this effect does not seem to be due to changes in the strength of PRDM9 binding to hotspots or in its multimer formation, as judged by *in vitro* experiments.

### Loss of EWSR1 affects DSB repair

To test whether EWSR1 influences recombination processes subsequent to PRDM9-dependent hotspot activation, we determined the numbers, locations, and activities of DSBs, marked by the presence of the protein DMC1 covering the single stranded DNA tails at DSBs [38], in CKO and control spermatocytes. We observed no significant difference in DMC1 foci number per cell between CKO and controls in early leptotene and zygotene spermatocytes (Fig. 4A).This result suggests that the *number* of DSBs per meiosis is not affected by the presence or absence of EWSR1.

**Fig 4.**
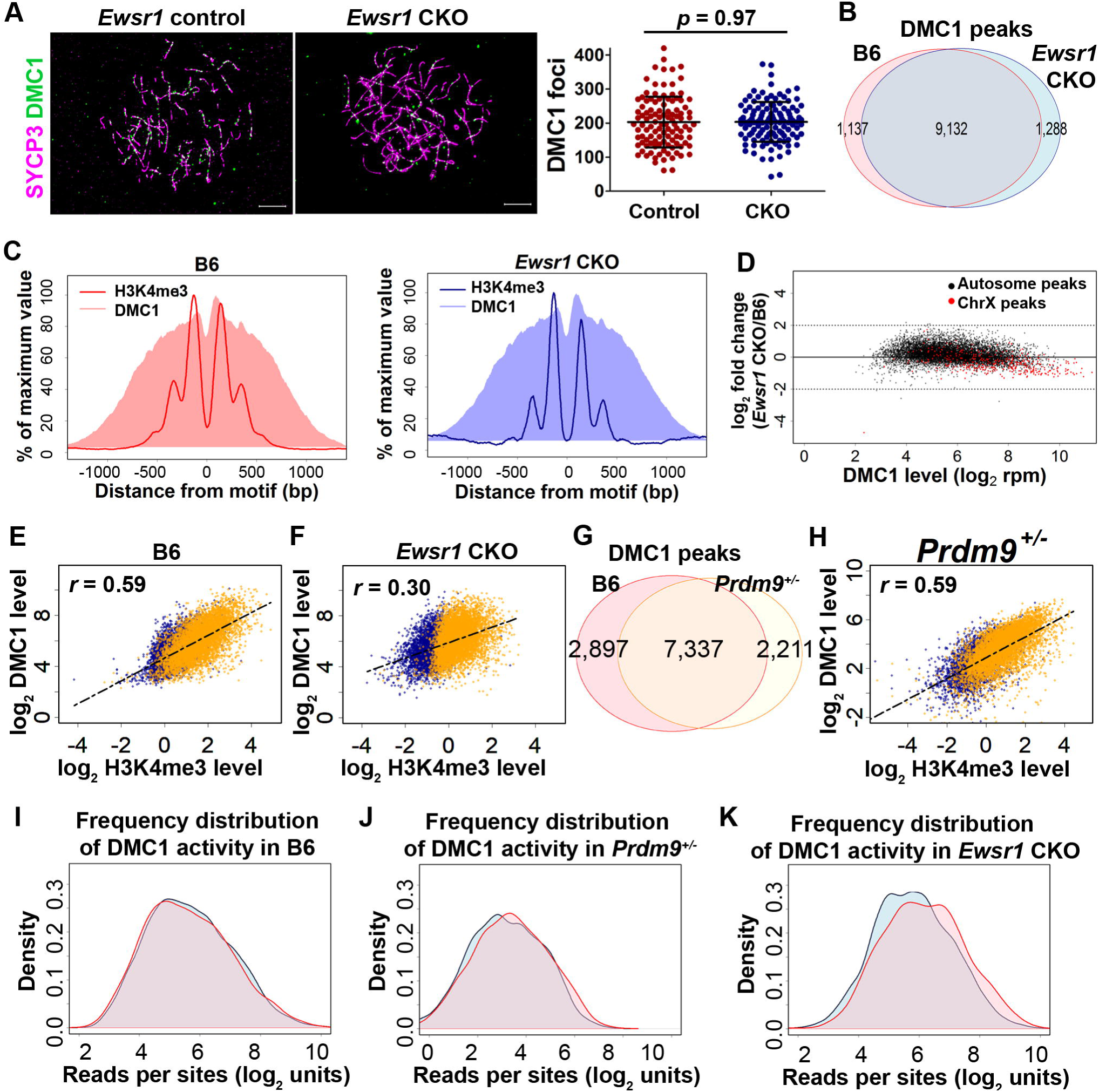
EWSR1 does not affect the total number of DSBs per cell but affects their positioning and activities. (A) Left - DMC1 staining on *Ewsr1* control and CKO chromosome spread. Scale bar, 10µm. Right – distribution plot of DMC1 foci in leptotene and zygotene spermatocytes. Bars represent mean ± SD. *p* = 0.96 by Student’s *t*-test. (B) DMC1 peaks occur at the same sites in B6 control (pink) and CKO (blue) spermatocytes. (C) Aggregation plot of H3K4me3 and DMC1 signal in B6 control (left) and CKO (right). The signal was normalized to the maximum signal. (D) MA-plot of activity of DSBs on autosomes (black dots, n = 8,744) and X chromosome (red dots, n = 388) from *Ewsr1* CKO and control spermatocytes. (E and F) Correlation between DMC1 level and H3K4me3 level in B6 (E) and *Ewsr1* CKO (F). Yellow dots, hotspots with significant H3K4me3 peaks in both control and CKO (FDR < 0.01). Blue dots, hotspots with significant H3K4me3 peaks in controls but not significant in CKO (FDR > 0.01). (G) Dosage effect of PRDM9 on hotspot initiation. Venn diagram of the DMC1 peaks identified from DMC1 ChIP-seq in control (pink) and *Prdm9^+/-^* (yellow) spermatocytes. (H) Correlation between DMC1 level and H3K4me3 level in *Prdm9^+/-^*. Yellow dots, hotspots with significant H3K4me3 peaks in both control and CKO (FDR < 0.01). Blue dots, hotspots with significant H3K4me3 peaks in controls but not significant in CKO (FDR > 0.01). (I, J and K) Frequency distribution of DMC1 activity in B6 (I), *Prdm9^+/-^* (J) and CKO (K) at hotspots in open (blue) and closed (red) chromatin.

To determine whether the *locations* or repair of DSB sites in *Ewsr1* CKO spermatocytes are affected, we first performed ChIP-seq for DMC1 [39, 40] in two replicates, which showed excellent correlation (Fig. S4A). We then asked whether DSBs still occur at PRDM9-dependent sites in the absence of EWSR1. We detected 10,420 DMC1 peaks in CKO spermatocytes, 87.6% of which (9,132 peaks) overlapped with hotspots in controls (Fig. 4B). Of those shared DMC1 peaks, 94.6% (8639 peaks) contained a PRDM9 binding site at their centers, and 78.0% of the shared peaks (7,122 peaks) overlapped H3K4me3 peaks, indicating that a great majority of DSBs in CKO spermatocytes were indeed initiated at PRDM9 binding sites. The aggregation plots also confirmed that DSBs contain PRDM9 binding motifs at their centers in both control and CKO spermatocytes (Fig. 4C).

Next, we asked whether the lack of EWSR1 affects DSB repair. In both CKO and control spermatocytes, DSBs on the X chromosome showed higher DMC1 signal compared to the autosomes (Fig. S4B), probably reflecting the delay of the DSB repair on the X chromosome until mid-pachytene [41]. On autosomes, the normalized overall activity of DMC1 in CKO spermatocytes was increased compared to controls (*p* = 9.6 × 10^-23^, Wilcoxon rank sum test), with a concomitant reduction on the X chromosome (*p* = 1.1 × 10^-41^, Wilcoxon rank sum test, Fig. 4D). This suggests that in CKO spermatocytes DSB repair is delayed on autosomes as well as on the X chromosome, which can explain the increased asynapsis we observed in CKO spermatocytes (Fig. 2F).

Together, these data suggest that in the absence of EWSR1 DSBs are still formed at PRDM9-dependent hotspots, but their repair is delayed.

### DSB activity and distribution are altered in the absence of EWSR1

Next, we investigated whether the loss of *Ewsr1* affects the positions and activities of DSB hotspots. In wild type spermatocytes, DMC1 levels generally correlate with H3K4me3 level (*r* = 0.59, *p* < 2.2 × 10^-16^, Fig. 4E). However, among the H3K4me3 hotspots in CKO spermatocytes, many of the weak hotspots, not detected at FDR < 0.01 (Fig. 4F blue dots), show similar DMC1 levels as the stronger hotspots (Fig. 4F yellow dots), resulting in a weaker correlation between H3K4me3 and DMC1 levels in CKO spermatocytes (*r* = 0.30, *p* < 2.2 × 10^-16^, Fig. 4F).To investigate whether this is the results of the reduced PRDM9 methyltransferase activity we observed in CKO spermatocytes (Fig. 3G), we carried out DMC1 ChIP-seq in *Prdm9^+/-^* spermatocytes which show reduction of H3K4me3 levels at hotspots similar to *Ewsr1* CKO mice (Fig. 3G, Fig.S4C and [35]). We detected 9,548 DMC1 peaks, of which only 76.8% (7,337 peaks) overlapped DMC1 sites in controls (Fig. 4G, correlation between the two replicas *r*=0.99, Fig. S4D), a significant difference compared to 87.6% overlap between *Ewsr1* CKO spermatocytes and controls (Fig. 4B). Of the remaining 2,211 non-overlapping peaks in *Prdm9^+/-^* spermatocytes, 1,606 (16.8% of the total of 9,548) did not contain a PRMD9 binding motif at the center. However, the correlation between DMC1 and H3K4me3 level in *Prdm9^+/-^* spermatocytes was as strong as the correlation in the controls (*r* = 0.59, *p* < 2.2 × 10^-16^; Fig. 4H, see also Fig. 4E). This discrepancy led us to test whether the redistribution of DSBs in the absence of EWSR1 is related to chromatin state [20]. In B6 and in *Prdm9^+/-^* spermatocytes, DMC1 levels were similar in open and closed chromatin (*p* = 0.21 in B6, *p* = 0.10 in *Prdm9^+/-^*, Wilcoxon rank sum test, Fig. 4I and 4J). In contrast, DSB activity was shifted towards closed chromatin in CKO spermatocytes (*p* = 1.21 × 10^-12^, Wilcoxon rank sum test; Fig. 4K and S4E).

Together, these results indicate that the reduced PRDM9 methyltransferase activity alone cannot explain the altered hotspot choice observed in *Ewsr1* CKO spermatocytes. They also suggest that one aspect of EWSR1 function in early meiotic prophase is participation in the selection of H3K4me3-activated hotspots for DSB formation.

### Lack of *Ewsr1* affects the number and distribution of genetic crossovers in meiosis

We tested whether the redistribution of DSBs caused by the loss of EWSR1 also affects the subsequent recombination events by determining the number and distribution of crossover sites, marked by the protein MLH1 [42] (Fig. 5A). Crossover number was significantly lower in CKO spermatocytes (18.3 ± 2.6) compared to control spermatocytes (22.9 ± 1.4; *p* <0.001) (Fig. 5B). This is less than the one crossover per chromosome that is required for normal meiosis. Indeed, at pachynema, about 70% of CKO spermatocytes lacked a crossover on the XY bivalent (yellow arrow in Fig. 5A, right panel, and Fig. S5A), and more than half had no crossover on at least one autosome (white arrow in Fig. 5A, right panel, and Fig. S5B).

**Fig 5.**
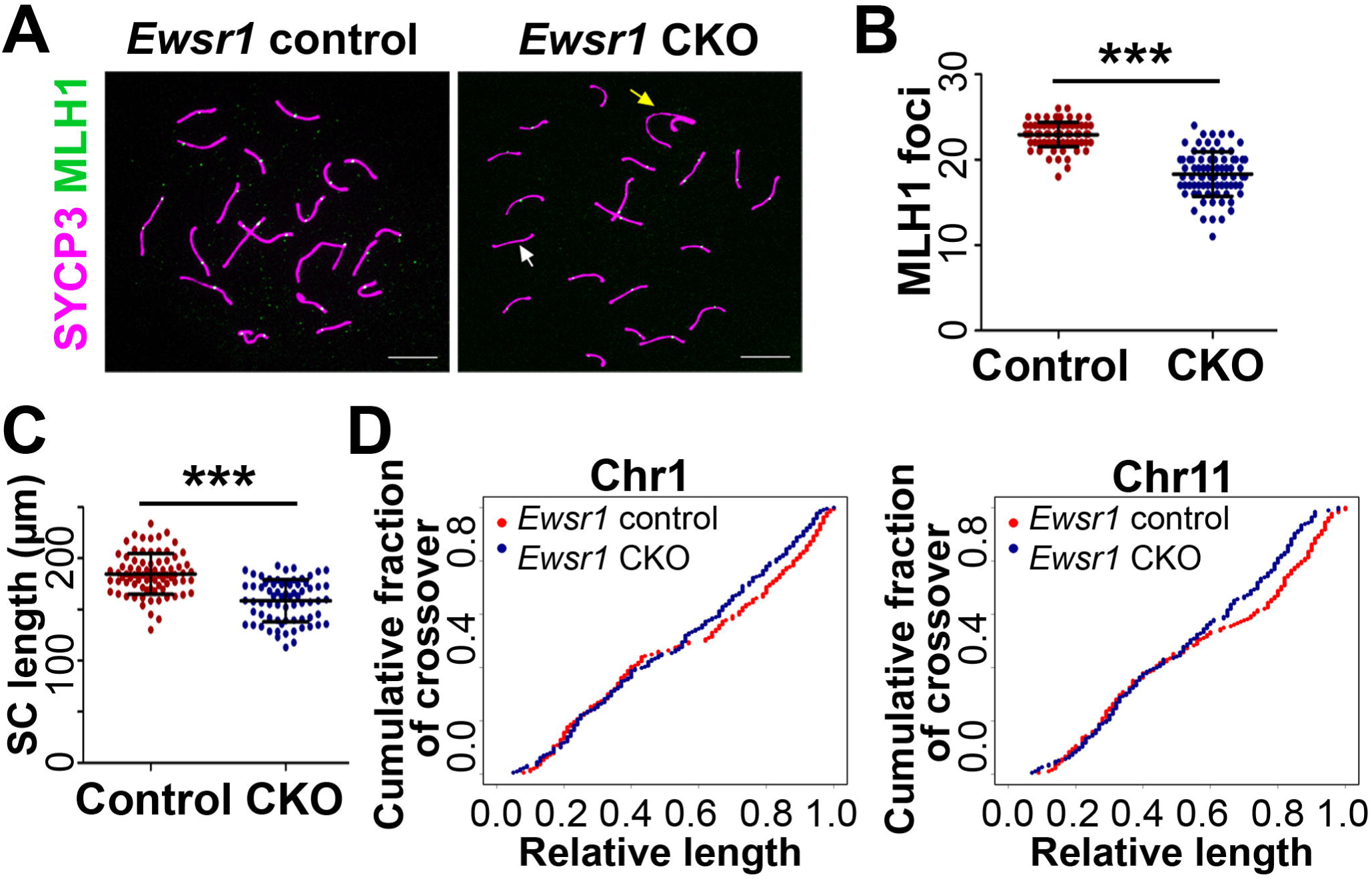
Crossover number and length of SC are reduced in *Ewsr1* CKO. (A) Presence of bivalents without crossovers in *Ewsr1* CKO. MLH1 (green) and SYCP3 (magenta) staining in *Ewsr1* control and CKO chromosome spread. White arrow, autosome with no crossover; yellow arrow, sex chromosomes with no crossover. Scale bar, 10 µm. (B) Distribution of the number of MLH1 foci in pachytene spermatocytes. (C) Distribution of the synaptonemal complex lengths. (D) Cumulative plots of crossovers on chromosome 1 (left) and chromosome 11 (right). Bars in (C) and (D) are mean ± SD. \*\*\**p* < 0.001 by Student’s *t*-test.

In human and mouse meiosis, synaptonemal complex (SC) length is correlated with crossover number [43–47]. Indeed, we observed a shorter SC length at pachynema in CKO spermatocytes (158.4 ± 20.4 µm) compared with *Ewsr1* control spermatocytes (184.7 ± 20.0 µm) (Fig. 5C).

The reduction in the total number of crossovers in CKO spermatocytes could be due to alteration of interference distance, as genetic interference is correlated with SC length [47], but this proved not to be the case. Instead, the reduction in the number of MLH1 foci was related to crossover repositioning along the chromosomes. The decrease in MLH1 foci numbers in *Ewsr1* CKO compared to control spermatocytes occurred in a similar pattern in long and middle-sized autosomes as illustrated by Chromosomes 1 and 11 (Fig. S5C and S5D). In both chromosomes, the number of bivalents with more than one MLH1 focus was smaller in CKO than in control spermatocytes (Fig. S5D). However, this reduction was not caused by a longer interference distance in CKO spermatocytes, as this distance—measured by the average SC distance between two foci on the same bivalent [48]—was similar in CKO and control spermatocytes, for each of the two chromosomes (Fig. S5E). We also examined the impact of EWSR1 on crossover distribution. MLH1-positive cells with double crossovers on Chromosome 1 or 11 showed highly similar double crossover distribution patterns in CKO and control spermatocytes (Fig. S5F top panels). In spermatocytes with single crossovers on Chromosome 1 or 11, the crossovers were more evenly distributed along the chromosomes in CKO compared to control spermatocytes in which crossovers are preferentially formed near the telomeres with a second frequency peak in the centromere-proximal third of the bivalents (Fig. 5D and Fig. S5F bottom panels). This redistribution of crossovers in CKO increases the likelihood of single crossovers and decreases the likelihood of two or more crossovers per bivalent.

Together, these results suggest that the absence of EWSR1 in early meiotic prophase affects downstream steps in the recombination pathway, including the choice of which H3K4me3-marked recombination hotspots will undergo DSBs and will ultimately be resolved as crossovers.

## Discussion

The results of this study indicate that EWSR1 is likely to be an important protein in early meiotic prophase whose action influences all subsequent recombination events, including the imposition of PRDM9-dependent H3K4/K36 trimethylation marks activating recombination hotspots, their subsequent usage for DSB formation, and the choice of DSBs destined for the crossover formation. We show that it does so by affecting two major processes: PRDM9-dependent H3K4/K36 trimethylation; and PRDM9-hotspot complex association with the nascent chromosome axis through pREC8. Although we show that EWSR1 participates in hotspot association with the chromosome axis, our data suggest that it cannot be the only protein to do so, because a significant share of PRDM9-dependent hotspots are forming DSBs in CKO spermatocytes. Another proposed candidate, CXXC1 [27, 49], does not seem to be important in the PRDM9-dependent hotspot recognition pathway [50], which raises the question of what other proteins, if any, may be involved in this process. In somatic cells, EWSR1 is reported to be involved in RNA-related functions such as transcription regulation and RNA binding [51, 52]. However, changes in transcription levels were not detected in the systemic *Ewsr1* mutant [28], and we did not find binding of EWSR1 to either mRNA or miRNA in meiotic cells (data not shown), indicating that those reported RNA-related functions of EWSR1 are probably not present in meiotic cells.

### EWSR1 enhances PRDM9 methyltransferase activity *in vivo*

In humans and mice, PRDM9 is the only known protein that determines the positions of meiotic hotspots [4–6] through its binding to specific DNA sequences and trimethylation of H3K4 and H3K36 at the adjacent nucleosomes [7, 14, 40]. Loss of PRDM9 [27, 29, 32, 39] or any of its functional domains—zinc fingers for DNA binding [27], PR/SET for methyltransferase activity, or KRAB domain for protein interactions [53]—result in germ cell apoptosis due to meiotic arrest. Here we found that EWSR1 influences the methyltransferase activity of PRDM9. Lack of EWSR1 causes two-to four-fold reduction of both H3K4 and H3K36 methyltransferase activities at all PRDM9 activity levels (Fig. 3D, 3E). This observation is not likely due to H3K4/36 methyltransferase activity of EWSR1, as EWSR1 has no histone modification-related domain [51]. EWSR1 could serve as activator of the PR/SET domain; however, this is unlikely, as purified full-length PRDM9 is fully capable of performing its two methyltransferase activities *in vitro* [7].

One plausible explanation is that loss of EWSR1 may affect PRDM9 methyltransferase activity by shifting the balance of PRDM9 binding to other proteins in the process of its interaction with hotspot DNA and/or with adjacent nucleosomes during meiotic initiation. We and others have identified several PRDM9 interaction proteins in addition to EWSR1, including histone modification readers or writers, such as histone acetyl reader CDYL, and H3K9 methyltransferase EHMT2 [27], interacting with PRDM9 through its KRAB and adjacent domains [27]. CDYL and EHMT2, both of which bind to PRDM9 through a region on PRDM9 overlapping the SET domain [27], may act as inhibitors of PRDM9 H3K4 methyltransferase activity; in somatic tissues, these two proteins form a complex that can methylate H3K9 and serve as a transcription repressor by imposing a closed chromatin mark [54]. Their expression pattern in spermatocytes suggests that they may be replaced as PRDM9 partners by EWSR1 and/or CHAF1A [27].Therefore, EWSR1 binding to PRDM9 may be necessary to release the activity of its SET domain *in vivo*. These various possible mechanisms through which EWSR1 could impact PRDM9 methyltransferase activity are not mutually exclusive.

### EWSR1 interacts with meiosis-specific cohesin pREC8 during meiosis

During meiosis, EWSR1 interacts not only with PRDM9 but also with the phosphorylated form of the meiosis specific cohesin REC8. This interaction is independent of the presence or absence of PRDM9 (Fig. 1 and [27]). Conversely, PRDM9 association with pREC8 and synaptonemal complex proteins is dependent on its binding to hotspot DNA [27]. Here we show that EWSR1 is necessary for PRDM9-hotspot DNA association with the cohesin elements of the chromosome axis, providing direct links to pREC8. EWSR1 does not seem to interact with the other meiosis specific kleisin cohesin subunit, RAD21L (Fig. 1A). Loss of PRDM9’s DNA binding, but not its methyltranferase activity, abolishes the association between PRDM9 and EWSR1 in spermatocytes (Fig. 6B and [27]), which suggests that PRDM9 binding to DNA is necessary for stabilization of the PRDM9-EWSR1 complex.

Our results suggest that the two EWSR1 functions in meiosis—enhancing PRDM9-dependent H3K4/K36 trimethylation and providing a link between PRDM9 and pREC8— may not be independent from each other. It is plausible that linking PRDM9-bound hotspots to the chromosome axis stabilizes the entire complex and allows effective H3K4/K36 trimethylation.

In yeast, Rec8 is required for normal DSB distribution and repair [55, 56]. In mice, loss of REC8 causes meiotic arrest, asynapsis and DSBs formation defects [57–59]. Moreover, in yeast and mammals, pREC8 is required for REC8 cleavage and meiotic prophase I progress [60–63], and is essential for crossover determination and SC formation [64]. Consequently, the EWSR1-dependent association between PRDM9-activated hotspots and pREC8 may promote proper DSB formation and crossover pathway choice, which can explain the altered DSB distribution, increased chromosomal asynapsis, and changes in crossover number and distribution that we observed in *Ewsr1* CKO spermatocytes.

### The EWSR1-PRDM9 complex affects the fate of subsequent recombination events

In *Ewsr1* CKO spermatocytes, although hotspot H3K4me3 number and level are remarkably reduced (Fig. 3D) and the association between PRDM9 and chromosome axis elements is impaired (Fig. 1D), the number of DSB per cell is not significantly different compared to controls (Fig. 4A). This is consistent with the presence of a homeostatic mechanism regulating the numbers of recombination events at several levels [16] and our previous observation that the number of DSBs is independent of *Prdm9* dose or hotspot trimethylation level [32]. Importantly, DSBs do not occur at PRDM9 binding sites in *Prdm9^Set-/Set-^* mice [19], confirming that PRDM9 binding is not sufficient to initiate DSBs at hotspots and that DSB initiation also requires the H3K4/K36 trimethylation activity of PRDM9. In the absence of EWSR1, the positions of DSBs are not significantly correlated with H3K4me3 levels (Fig. 4F). However, we found that the distribution of H3K4me3 signals at hotspots as well as at non-hotspots is not affected in *Ewsr1* CKO spermatocytes. Specifically, in wild type controls we found that only 10.90% of hotspot peaks and 5.84% of non-hotspot peaks are detected within the published H3K9me3 domains, which contribute to 10.57% and 3.69% of total hotspot and non-hotspot activity, respectively. In *Ewsr1* CKO spermatocytes, 7.24% of hotspot peaks and 4.52% of non-hotspot peaks are detected within the H3K9me3 domains, which contribute to 7.14% and 2.99% of total hotspot and non-hotspot activity, respectively.

Further, EWSR1 has no reported histone modifying domain [51] and PRDM9 is not involved in H3K9me2/3 activities *in vitro* [20]. Therefore, it is unlikely that the DSB activity shift to closed chromatin is caused by EWSR1 or PRDM9 affecting the open or closed chromatin states directly. This suggests that EWSR1 is actively involved in associating hotspots with specific cohesin complexes forming the chromosome axis and thereby promotes recombination [26], although it is possible that EWSR1 isnot the only protein to do so. For example, in the absence of EWSR1, CXXC1 could link PRDM9-trimethylated hotspots to another chromosome axis element, IHO1, which is not a cohesin [53]. This alternative pathway could explain the shift of DSB activity along the chromosomes and towards closed chromatin (Fig. 4K) in CKO spermatocytes, which could reflect either a true increase in DSB frequency in closed chromatin, or, more likely, initiation of equal numbers of DSBs in open and closed chromatin, but with a longer period of time required for repair of DSBs in closed chromatin.

The shifted pattern of DSB initiation towards closed chromatin and away from telomeres (Fig. 4K) affects the subsequent recombination repair pathways and results in reduced number of crossovers to less than one crossover per bivalent (Fig. 5B), thereby breaking the crossover assurance efficiency [65–68]. This reduced number is not a result from increase in crossover interference; it reflects the more random crossover positioning in CKO losing the typical wild type pattern of bimodal distribution of crossovers relative to the centromere-telomere axis (Fig. 5B) [69–71]. As a result of this more random positioning, the fraction of bivalents with double or triple crossovers (Fig. S5D), as well as the SC length (Fig. 5C), are reduced in CKO spermatocytes; the latter is known to correlate strongly with DSB and crossover number [43, 72].

The reduced SC length in *Ewsr1* CKO spermatocytes is reminiscent of that of *Rec8* mutant [57, 73, 74], further suggesting that the two proteins act in the same recombination pathway. In the *Ewsr1* CKO spermatocytes, both SC length and crossover number are reduced, which indicates that loss of EWSR1 alters DSB repair pathways most likely through disassociation between hotspots and pREC8.

In summary, our results suggest that EWSR1 has a dual action in early meiotic prophase. In leptonema, EWSR1 joins the recombination-initiating complex of PRDM9 dimer/oligomer – hotspot DNA forms (and simultaneously binds to pREC8, incorporating the recombination initiation complex into the nascent chromosome axis and resulting in more efficient H3K4/K36 trimethylation at the nearby nucleosomes. The preferential link to meiotic-specific cohesin complexes containing pREC8 promotes the crossover pathway of DSB repair [12, 13] and ensures correct positioning of crossovers relative to the centromere-telomere axis. In the absence of EWSR1, the recombination initiation complexes are not actively associated with pREC8. This results in shifted positioning of crossovers relative to the centromere-telomere axis and disruption of crossover assurance, leaving some bivalents without crossovers and causing germ cell apoptosis in late prophase I.

## Materials and Methods

### Ethics Statement

The animal care rules used by The Jackson Laboratory are compatible with the regulations and standards of the U.S. Department of Agriculture and the National Institutes of Health. The protocols used in this study were approved by the Animal Care and Use Committee of The Jackson Laboratory (Summary #04008). Euthanasia for this study was done by cervical dislocation.

### Mice

All wild-type mice used in this study were in the C57BL/6J (B6) background. The mouse line C57BL/6N-Ewsr1^<tm1c(EUCOMM)Wtsi>^/Tcp was generated as part of the NorCOMM2 project at the Toronto Centre for Phenogenomics [75], and obtained from the Canadian Mouse Mutant Repository. The *Ewsr1* conditional knockout mice used in this study were produced by a two-step deletion scheme. C57BL/6N-Ewsr1^<tm1c(EUCOMM)Wtsi>^/Tcp mice were firstly backcrossed to C57BL/6J, and then mated to Tg(Sox2-cre)1Amc/J mice to generate one *Ewsr1* allele deleted mice, and then the *Ewsr1* hemizygous mice (*Ewsr1*^Δ/+^) were mated to Tg(Stra8-icre)1Reb/J to obtain *Ewsr1*^Δ/+;Stra8-iCre^ mice. The *Ewsr1*^Δ/+;Stra8-iCre^ mice were mated to homozygous Ewsr1 loxP mice to generate heterozygous control mice (*Ewsr1^loxp/+;Stra8-iCre^*, designated as Ewsr1 control) or conditional knockout mice (*Ewsr1^loxp/+;Stra8-iCre^*, designated as *Ewsr1* CKO).

B6(Cg)-Prdm9tm2.1Kpgn/Kpgn mice (designated as *Prdm9^Set-/Set-^*) were generated by introducing a point mutation creating G274A amino acid substitution that causes a total loss of PRDM9 PR/SET domain methylation activity. B6(Cg)-Prdm9tm3.1Kpgn/Kpgn mice (designated as *Prdm9*^ΔZnF/ΔZnF^) were generated in a previous study [27]. B6;129P2-Prdm9^tm1Ymat^/J mice (designated as *Prdm9^-/-^*) have been previously described [29]. All animal experiments were approved by the Animal Care and Use Committee of The Jackson Laboratory (Summary #04008).

### Measurement of testis index

Testicular weight and body weight of 14 dpp *Ewsr1* control (n = 3) and CKO (n = 4), 18 dpp *Ewsr1* control (n = 6) and CKO (n = 6) and 8 weeks *Ewsr1* control (n = 6) and CKO (n = 6) mice were measured. Testis index was calculated as testis weight/body weight. T-test was used to determine the statistical significance.

### Fertility test

Fertility test was performed with 4 *Ewsr1* control and 3 CKO male mice. Each mouse mated with at least two B6 females for at least three month period. Litter size and viable pup number were record.

### Histology

Testis or epididymis from adult B6, *Prdm9^Set-/Set-^*, *Ewsr1* control or CKO mice were dissected out, fixed with Bouin’s solution, and embedded in paraffin wax, and 5-μm sections were prepared. Sections of testis were stained with Periodic acid–Schiff– diastase (PAS), and section of epididymis were stained with haematoxylin and eosin (HE) using standard techniques. The slides were scanned by Nanozoomer.

### TUNEL assay

For detection of apoptosis in tissues, testis sections from *Ewsr1* control (n = 3) and CKO (n = 3) were subjected to fluorescence labeling of DSBs by terminal deoxynucleotidyl transferase–mediated digoxigenin-dUTP nick-end labeling (TUNEL) assay, using the In Situ Cell Death Detection Kit (11684795910, Roche) according to the manufacturer’s protocol. The images were captured by Zeiss imager Z2 microscope. The TUNEL positive cell number per seminiferous tubule was counted. T-test was used to determine the statistical significance.

### Chromosome spread and FISH

The drying-down technique [76] was used for preparation of chromosome spreads from spermatocytes. Briefly, testes from 8-weeks B6, *Prdm9^Set-/Set-^*, *Ewsr1* control or CKO mice were removed and decapsulated. The seminiferous tubules were incubated in hypotonic extraction buffer (30 mM Tris-HCl pH 8.3, 50 mM sucrose, 17 mM sodium citrate, 5 mM EDTA, 5 mM DTT) with 1X phenylmethanesulfonylfluoride (PMSF) for 30 min, transferred to 100 mM sucrose and disrupted using two fine forceps. Cell suspension was dropped on a slide covered with 1% paraformaldehyde solution. The slides were dried overnight in a humid chamber at room temperature. The slides were then washed with 0.4% Photo-Flo and blocked with 1X ADB (0.3% BSA, 1% normal donkey serum and 0.05% Triton X-100), following by immunolabeling with anti-PRDM9 (1:200), SYCP3 (1:400), γH2AX (1:1000, Abcam, ab26350), BRCA1 (1:200, Santa Cruz, sc-642), ATR (1:500, Santa Cruz, sc-1887), HORMAD1 (1:500, Protein Tech, 13917-1-AP), H3K4me3 (1:1000, 07-473, Millipore), H1t (1:1000, custom-made), DMC1 (1:200, Santa Cruz, sc-8973) or MLH1 (1:100, BD Pharmingen, 550838) antibodies. The images were captured by Zeiss imager Z2 microscope. The numbers of cells in different spermatocyte stages were counted using chromosomes spreads with SYCP1/SYCP3/histone variant H1t (marks late pachytene cells) staining; *Chi*-square was used to determine the statistical significance. The number of DSBs was counted by DMC1 foci per leptotene/early zygotene spermatocytes in *Ewsr1* control (n = 111, from three individual mice) and CKO (n = 120, from three individual mice). The number of crossover was counted by MLH1 foci per pachytene spermatocytes in *Ewsr1* control (n = 86, from three individual mice) and CKO (n = 81, from three individual mice). The SC length was measure as SYCP3 length of pachytene spermatocytes in *Ewsr1* control (n = 78, from three individual mice) and CKO (n = 70, from three individual mice). T-test was used to determine the statistical significance.

After MLH1/SYCP3 immunostaining on chromosome spread, images were captured and analyzed, and fluorescence in situ hybridization (FISH) was carried out on the same spermatocytes using chromosome 1-(MetaSystems, XMP1 Green) and chromosome 11-(MetaSystems, XMP11 Orange) specific probes according to the manufacturer’s protocol. Each spermatocyte was re-captured to identify chromosome 1 and 11, and the distance of each crossover to centromere on those two chromosomes was measure in 118 *Ewsr1* control pachytene spermatocytes from 2 mice and 178 CKO pachytene spermatocytes from 2 mice. Student’s *t*-test and *Chi*-square was used to determine the statistical significance.

### Immunofluorescence

For protein immunolocalization on tissue sections, our established method was used [27]. Briefly, tissues from 8 week old *Ewsr1* control and CKO were dissected out, fixed with 4% paraformaldehyde solution overnight, embedded in paraffin wax, and sectioned at 5 μm. Sections were heated in a microwave in 10 mM sodium citrate buffer, pH 6.0, for 10 min and treated with phosphate-buffered saline (PBS) containing 0.1% Triton X-100. After blocking of nonspecific binding sites with 10% normal donkey serum, sections were stained with antibodies against EWSR1 (1:200, Abcam, ab54708), PRDM9 (1:200, custom-made), EHMT2 (1:1000, Cosmo Bio Co, PP-A8620A-00) and CDYL (1:50, Abcam, ab5188). The images were captured by Zeiss imager Z2 microscope.

### H3K4me3 and H3k36me3 ChIP-seq and analysis

H3K4me3 and H3k36me3 ChIP was performed in two replicates with 14 dpp B6 and *Ewsr1* CKO spermatocytes, and one replicate with 14 dpp *Prdm9^Set-/Set-^* spermatocytes, using the protocol we reported previously [14]. Briefly, spermatocytes were isolated and crosslinked with 1% formaldehyde solution for 10 min at room temperature. After washed with PBS and spun down, spermatocytes were incubated in 1 ml hypotonic lysis buffer (10 mM Tris-HCl pH 8.0, 1 mM KCl, 1.5 mM MgCl_2_) with 1 mM PMSF and 1× EDTA-free protease inhibitor cocktail (PIC, Roche, 11873580001) for 30 min at 4°C. Nuclei were pelleted by spinning down at 10,000 x *g* for 10 min and resuspended by MNase buffer (50 mM Tris-HCl pH 8.0, 1 mM CaCl, 4 mM MgCl, 4% NP-40) with 1 mM PMSF and 1×PIC. 15 U per 5X10^6^ spermatocytes MNase was added into the cell mixture to fragment chromatin, followed by 8 min incubation at 37°C. The reaction was stopped by adding 10 mM EDTA. Chromatin was clarified by centrifugation at top speed to spin down the insoluble fragment at 4°C for 10 min. The supernatant was transferred to a new tube, and 25% of volume was saved as input chromatin. Chromatin was incubated with protein A Dynabeads (Thermo Fisher Scientific, 10002D) conjugated with antibodies against H3K4me3 (Millipore, 07-473) or H3K36me3 (Active Motif, 61101) overnight at 4°C. After incubation, the beads were washed with 1 ml RIPA buffer for three times and TE buffer for twice. DNA was eluted with 125 μl elution buffer (1% SDS, 20 mM Tris-HCL pH 8.0, 200 mM NaCl, 5 mM EDTA) supplemented with 50 μg/ml Proteinase K (Sigma-Aldrich, 39450-01-6) and reverse-crosslinked at 68°C for 2 hrs with vigorous shaking at 14,000 rpm in a thermal mixer. ChIP DNA was recovered from the beads using magnetic separation, and purified with QIAquick PCR Purification Kit (Qiagen, 28106) following the manufacturer’s protocol. Libraries preparation for sequencing was performed by using NEXTflex ChIP-Seq Kit (Bioo Scientific, NOVA-5143-01) without size selection.

ChIP libraries were sequenced on an Illumina^®^ HiSeq 2500 platform, with 75 bp single-end reads. H3K4me3 and H3k36me3 ChIP reads were trimmed for quality using trimmomatic (v.0.32), and then aligned to the mouse mm10 genome using BWA (v.0.5.10-tpx). Data are available at NCBI Gene Expression Omnibus (GEO; http://www.ncbi.nlm.nih.gov/geo) under accession number GSE108259 (Accession for review purpose: https://www.ncbi.nlm.nih.gov/geo/query/acc.cgi?acc=GSE108259). Peak calling was performed using MACS (v.1.4.2) with standard treatment (ChIP) and control (input) samples with a false discovery rate (FDR) value 0.01. H3K4me3 activity was normalized in two methods: (1) to reads per million (rpm); (2) to a set of promoters (*Mlh1, Prdm9, Pms2, Mnd1, Dmc1, Rad21l, Mei4, Stra8, Ctcfl, Mei1, Puf60, Eef2, Rpl38, Leng8, Setx, Eif3f, Rpl37, Psmd4, Heatr3, Chmp2a, Sycp1, Sycp2, Sycp3, Morc2b, Zfp541, Spo11, Mdh1b, Rec8, Msh4, Psmc3ip*) used to normalize H3k4me3 enrichment in previous studies [19, 77]. The two methods resulted in similar normalization factors for the *Ewsr1* CKO sample (1.1560 in method 1 and 1.3298 in method 2). Therefore, we used reads per million for normalization in the following analysis. H3K36me3 activity was normalized to reads per million (rpm). PRDM9^Dom2^ hotspot locations were previously described by [20] (GSE52628) and lifted over to the mouse genome version mm10. Open or closed chromatin regions in spermatocytes were determined by H3K9me3 domains, which were previously described by [20] (GSE61613) and also lifted over to the mouse mm10 genome. H3K4me3 peaks shared by B6, *Prdm9^Set-/Set-^* and CKO spermatocytes, inside and outside known hotspots, in open/closed chromatin regions or H3K36me3 peaks in genic/intergenic regions were determined using bedtools (v2.22.0) intersect. Wilcoxon rank sum test was used to calculate the statistical significance. Analyses for the aggregation plots were carried out using the Aggregation and Correlation Toolbox (ACT) [78]. ACT parameters were: nbins = 500, mbins = 0, radius = 1500.

### DMC1 ChIP-seq and data analysis

DMC1 ChIP was performed with two replicates of 8 weeks old B6*,Prdm9^Set-/ Set-^*, *Prdm9^+/-^*and *Ewsr1* CKO spermatocytes using an established method [79]. Briefly, testes from 8-weeks mice were removed and decapsulated, and cross-liked with 1% paraformaldehyde solution for 10 min. The tissue was homogenized and filtered with 40 μm cell strainer to obtain germ cells. Cells were washed with Lysis buffer 1 (0.25% Triton X100, 10mM EDTA, 0.5mM EGTA, 10mM Tris-HCl pH 8.0), Lysis buffer 2 (0.2M NaCl, 1mM EDTA, 0.5mM EGTA, 10mM Tris-HCl pH 8.0), and resuspended in Lysis buffer 3 (1% SDS, 10mM EDTA, 50mM Tris-HCl pH 8.0) with 1X PIC. The chromatin was then sheared to ∼1000 bp by sonication, dialyzed against ChIP buffer (0.01% SDS, 1.1% Triton X-100, 1.2 mM EDTA, 16.7 mM Tris-HCl, pH 8.0, 167 mM NaCl) for at least 6 hrs. 2 µl of chromatin was saved as input. The rest of chromatin was incubated with antibody against DMC1 (Santa Cruz, sc-8973) overnight at 4°C. The mixture was then incubated with protein G Dynabeads (Thermo Fisher Scientific, 10004D) for 4 hrs at 4°C. The beads were washed with Wash buffer 1 (0.1% SDS, 1% Triton X-100, 2 mM EDTA, 20 mM Tris-HCl, pH 8.0, 150 mM NaCl), Wash buffer 2 (0.1% SDS, 1% Triton X-100, 2mM EDTA, 20mMTris-HCl, pH 8.0, 500 mM NaCl), Wash buffer 3 (0.25 M LiCl, 1% NP-40, 1mM EDTA, 10mMTris-HCl, pH 8.0, 1% Deoxycholic acid) and twice with TE buffer. The chromatin was eluted with dilution buffer (1% SDS, 0.1 M NaHCO_3_ pH 9.0) at 65°C for 30 min and reverse-crosslinked by adding 200 mM NaCl and incubation overnight at 65°C. ChIP and input DNA was purified by MinElute Reaction Cleanup Kit (Qiagen, 28006). For library preparation, DNA was firstly end repaired by incubation with 1X T4 DNA Ligase Reaction Buffer with 0.25 mM dNTP, 3 units of T4 DNA polymerase (NEB, M0203S), 1 unit of Klenow Enzyme (NEB, M0210S) and 10 units of T4 PNK (NEB, B0202S) at 20°C for 30 min, followed by addition of 3’-A overhangs using Klenow Fragment 3’-5’ exo^-^ (NEB, M0212S). After denaturation of DNA at 95°C for 3 min, the adapters from TruSeq Nano DNA LD Library Prep Kit (set A, Illumina, FC-121-4001) were ligated with Quick Ligation kit (NEB, M2200S). The libraries then were amplified using PCR Enhancer mix and primer cocktail (Illumina, FC-121-4001) for 12 cycles. For each step, DNA was purified by MinElute Reaction Cleanup Kit.

Libraries were sequenced on an Illumina® HiSeq 2500, with 75 bp paired-end reads. Fastq files for paired-end sequenced DMC1 samples were trimmed using Trimmomatic (v0.32) and subsequently parsed for detection and selection of paired reads having homology at the 5’ and 3’ ends as established by protocols for single strand DNA enrichment. The resultant files contain only the detectable single strand reads. These files were aligned to mm10 genome using BWA (v.0.5.10-tpx) and bam files were parsed for detection and selection of reads containing true genomic sequence versus fill-in sequence at the homologous region. These reads were selected from the original paired-end fastq files and single-ended fastq files were created containing only the true genomic sequences of single strand DNA reads.

The filtered reads were then aligned to the mouse mm10 genome using BWA (v.0.5.10-tpx). Data are available at NCBI Gene Expression Omnibus (GEO; http://www.ncbi.nlm.nih.gov/geo) under accession number GSE108259 (Accession for review purpose: https://www.ncbi.nlm.nih.gov/geo/query/acc.cgi?acc=GSE108259). 11,3240,405 and 13,474,644 aligned DMC1 reads in B6, 5,592,954 and 8,736,645 aligned DMC1 reads in *Ewsr1* CKO spermatocytes, 6,223,788 and 7,088,973 aligned DMC1 reads in Prdm9Set-/Set-spermatocytes, 5,137,175 and 4,413,135 aligned DMC1 reads in *Prdm9^+/-^* spermatocytes were obtained from each libraries. The correlation between the two biological replicates in each experiment was high (r = 0.95 in *Ewsr1* CKO; r = 0.99 in B6. Fig S5A); thus, the data from each pair of replicates were merged. The DMC1 activity was normalized to reads per million (rpm). Peak calling was performed using MACS (v.1.4.2) with standard treatment (ChIP) and control (input) samples with a FDR value 0.01. PRDM9^Dom2^ hotspot H3K9me3 domain locations were used same as in H3K4me3 ChIP-seq analysis. DMC1 peaks shared by B6, *Prdm9^Set-/Set-^*, *Prdm9^+/-^* and CKO spermatocytes, inside and outside known hotspots, or open/closed chromatin regions were determined using bedtools (v2.22.0) intersect. Wilcoxon rank sum test was used to calculate the statistical significance. Analyses for the aggregation plots were carried out using the ACT[78], of which parameters were: nbins = 500, mbins = 0, radius = 1500.

### Western blot

Testes from 14 dpp *Ewsr1* control or CKO mice were lysed in RIPA buffer with 1 mM PMSF and 1X PIC. The testicular extract was run on a SDS-PAGE 4-15% mini-protean TGX precast gel (Bio-Rad, 456-1084) and transferred to a nitrocellulose membrane. Anti-PRDM9 (1:1000, custom made), EWSR1 (1:1000, Abcam, ab54708), REC8 (1:1000, Abcam, ab38372), EHMT2 (1:1000, Cosmo Bio Co, PP-A8620A-00), SYCP3 (1:1000, Novus, NB300-231) antibodies were used for detecting proteins. anti-β-tubulin (1:1000, Sigma-Aldrich, T4026) was used as internal control.

### Protein purification and in vitro pull-down

6His-Halo-MBP-tagged and MBP-tagged PRDM9, and GST-tagged EWSR1, were expressed and purified as described previously [27]. Briefly, expression of those proteins was performed in Rosetta2 cells. Pre-culture was grown overnight at 30°C, and then the cells were re-inoculated in the next day and cultured for 7 hrs at 37°C. The cells were spun down and resuspended in 20 ml of Cell breakage buffer (50 mM Tris-HCl, 4 mM EDTA, 200 mM sucrose, pH 7.5) added with 0.1% NP-40, 0.1% β-cap, 100mM KCl and 1X PIC. After sonication to lysis the cells, the supernatant was purified by Sp-Sepharose.

The purified proteins were pre-cleared with MyOne T1 beads (Thermo Fisher Scientific, 65602) for 1.5 hrs. Then, HaloTag^®^ PEG-Biotin Ligand (Promega Corporation, G8591) was added to pre-cleared 6His-Halo-MBP PRDM9 to 1 μM final concentration and the mixture was incubated 2 hrs at room temperature. The pull down was then performed by mixing 1 μg of each protein into the mixture and incubating for 1 hr. The protein mixture was combined with MyOne T1 beads and incubated 1 hr at room temperature. The supernatant was saved. After washing with K buffer (40 mM K_2_PO_4_, 10% glycerol, 0.5 mM EDTA, pH 7.5), the bound proteins were eluted with SDS loading buffer. The samples were analyzed by SDS-PAGE and western blotting by using antibodies against MBP (1:1000, New England Biolabs, E8038S) or GST (1:1000, Sigma-Aldrich, GE27-4577-01).

### Electrophoretic mobility shift assay

Electrophoretic mobility shift assay (EMSA) was performed as previously described [37]. Two PRDM9^Dom2^ hotspots were selected for EMSA, including a strong hotspot Pbx1 and a moderate activated hotspot at Chr16 66.9 Mb. DNA fragment of those two hotspots were amplified by PCR from B6 genomic DNA, using the biotinylated primers (Eurofins MWG Operon). 0.6 ng of MBP-tagged PRDM9 and 0.7 ng of 6His-HALO-tagged EWSR1 were mixed in the EMSA buffer (10mM Tris-HCl, pH 7.5, 50mM KCl, 50 ng/μl poly(dI/dC) and 0.05% NP-40) for 20 min at room temperature. Total amount of proteins was adjusted by empty MBP or 6His-Halo tags to ensure that the protein concentrations in each reaction were equal. 20 ng 3’ biotin labeled oligo was added into the mixtures. The mixtures were incubated at room temperature for 20 min, and then separated on 5% PAA gel in 0.5× Tris-borate-EDTA (TBE) buffer. After electrophoretic, the products were transferred to nylon membrane by wet transfer in 0.5× TBE. The membrane was crosslinked by UV light at 120 mJ/cm^2^ and blocked with 10× EMSA blocking buffer (125 mM NaCl, 17 mM Na_2_HPO_4_, 8.3 mM NaH_2_PO_4_ and 5% SDS) for 30 min at room temperature. The membrane was then incubated with the streptavidin-peroxidase conjugate in 10× EMSA blocking buffer for 30 min at room temperature, and wash with 1× EMSA blocking and 1× washing buffer (10 mM Tris-HCl, pH 9.5, 10 mM NaCl, 20 mM MgCl_2_). The biotin-labeled products were detected by the chemiluminescence signal to the manufacturer’s specification. The density of band was measured by ImageJ (Fiji). The proportion of shifted band was calculated as S/(S+U), where S is the density of the shifted band and U is the density of the unshifted band. The experiment was done by three independent replicates. Two-way ANOVA was used to calculate the statistical significance.

### Co-immunoprecipitation assays

The co-IP was carried out using our reported protocol [27]. Testes of twenty 14 dpp B6, *Prdm9*^ΔZnF/ΔZnF^, *Prdm9^Set-/Set-^*, *Ewsr1* control or CKO mice were homogenized in cold PBS and passed through a 40-μm cell strainer (Falcon BD, 352340), and centrifuged for 5 min at 3000 × *g.* The pellet was resuspended and incubated for 30 min in 1 ml of Pierce IP buffer (ThermoFisher Scientific, 87787) with 1 mM PMSF and 1× PIC, and then span down at 13,200 × *g*. For the DNase I-treated co-IP samples, the supernatant was added with 100 μl of DNase I buffer and 20 U DNase I (Thermo Fisher Scientific, AM1906), and incubated for 1 hr at room temperature. After incubation, 10% of extract was set apart as input. The co-IP was perform by incubating extract with protein A or G Dynabeads conjugated with antibodies against PRDM9 (custom-made) or EWSR1 (sc-6532, Santa Cruz) overnight at 4°C. IgG from the same animal species was used as negative control. The beads were washed three times with 1 ml of Pierce IP buffer and eluted with 200 μl of GST buffer (0.2 M glycine, 0.1% SDS, 1% Tween 20, pH 2.2) for 20 min at room temperature. The sample was then neutralized with 40 μl of 1 M Tris-HCl, pH 8. After heated at 95°C for 5 min, 10 µg of IP and input samples were then subjected to electrophoresis and western blotting for detection of PRDM9 (1:1000, custom made), EWSR1 (1:1000, Abcam, ab54708), REC8 (1:1000, Abcam, ab38372), RAD21L (1:600, gift from Jibak Lee), SMC3 (1:1000, Abcam, ab9263), STAG1 (1:1000, Abcam, ab4455), SYCP3 (1:1000, Novus Biologicals, NB300-231) or SYCP1 (1:1000, Novus Biologicals, NB300-229).

## Acknowledgments

This work is dedicated to the memory of Pavlina M. Petkova, a talented scientist and a dear colleague. We thank Anita Adams and Catrina Spruce for technical help, Mary Ann Handel and Jibak Lee for sharing their antibodies, and Ken Paigen for critical reading and helpful suggestions. This work was supported by National Institutes of Health Grants R01 GM078452 to P.M.P., P50 GM076468 to GaryChurchill/Project B to P.M.P., and Cancer Core Grant CA34196 to the Jackson Laboratory.

## Supporting Information

**Fig S1 – related to Fig 1. EWSR1 interacts with meiotic cohesin protein pREC8.** Co-IP with EWSR1 from 14 dpp B6 testis extract. Staining for SYCP3 and SYCP1. In each blot, lane 1 – input; lane 2 – co-IP with non-immune IgG; lane 3 – co-IP with anti-EWSR1 untreated with DNAse I; lane 4 – co-IP with anti-EWSR1 treated with DNAse I.

**Fig S2 –related to Fig 2. Ewsr1 CKO mice are sterile with meiotic arrest and chromosomal asynapsis in spermatocytes.** (A) Testis index (testis weight/body weight) in 14-, 18-dpp and 8 weeks *Ewsr1* control (n = 3, 6 and 6, respectively) or CKO mice (n = 4, 6 and 6, respectively). (B) Percentage of late pachytene spermatocytes (marked by H1T) in *Ewsr1* control (n = 554) and CKO spermatocytes (n = 503). Spermatocyte stage proportion in adult *Ewsr1* control (983, 655 and 340 cells counted from 3 mice) and CKO mice (996, 652 and 196 cells counted from 3 mice). γH2AX (magenta) staining in different stage spermatocytes from *Ewsr1* control and CKO mice. Arrow, asynapsed regions. (E) ATR, BRCA1 and HORMAD1 (magenta) and SYCP3 (green) staining on *Ewsr1* control and CKO spermatocytes chromosome spreads. Arrows indicate asynapsed chromosomes in CKO. Scale bar, 10 µm. Bars in (B)-(D) represent mean ± SD. * *p* < 0.05, *** *p* < 0.001 by Student’s *t*-test.

**Fig S3 – related to Fig 3. H3K4me3 and H3K36me3 activities at hotspots are reduced in Ewsr1 CKO.** (A) Immunoblots of EWSR1, PRDM9 and PRDM9 interactors in Ewsr1 control and CKO testicular extract. (B) EHMT2 and CDYL staining in *Ewsr1* control and CKO seminiferous tubule sections. Scale bar, 50 µm. (C) H3K4me3 (green) and SYCP3 (magenta) staining on B6 and *Prdm9^-/-^* chromosome spreads in different stage spermatocytes. Scale bar, 10 µm. (D) H3K36me3 (green) and SYCP3 (magenta) staining on *Ewsr1* control and CKO. Scale bar, 10 µm. (E) Plots of two replicates of B6 controls or *Ewsr1* CKO H3K4me3 ChIP-seq samples. (F) Activity of two hotspots that were selected for EMSA. Black dot, a strong hotspot Pbx1; blue dot, a moderately activated hotspot at Chr16 66.9Mb. (G) EMSA with MBP-PRDM9 and DNA fragment of the hotspot at Chr16 66.9Mb in the presence or absence of 6His-HALO-EWSR1. Red arrow, shifted band; black arrow, unbound fragment. Right, quantification of shifted fraction. Data represent mean ± SD. *** *p* < 0.001, N.S., not significant, by two-way ANOVA. (H and I) Pull-down assay with purified 6HisHALO-MBP-PRDM9, MBP-PRDM9 (H and I) or empty MBP-tag (I). SN, supernatant. Beads, elution from beads.

**Fig S4 – related to Fig 4. EWSR1 does not affect the total number of DSBs per cell.** (A) Plots of two replicates of B6 controls or *Ewsr1* CKO DMC1 ChIP-seq samples. (B) Density plots of DMC1 activity in autosomes (grey) and X chromosome (red) in B6 (top) and CKO (bottom) spermatocytes. (C) Coverage profiles of H3K4me3 (blue) and DMC1 (red) from a representative 200-kb window in *Ewsr1* control, CKO, *Prdm9^Set-/Set-^* and *Prdm9^+/-^* spermatocytes. Red box, a PRDM9 dependent hotspot; blue box, promoter of Qk. (D) Plots of two replicates of *Prdm9^Set-/Set-^* DMC1 ChIP-seq samples. (E) MA-plot of DMC1 level in open (blue) and closed (red) chromatin of B6 and *Ewsr1* CKO.

**Fig S5 – related to Fig 5. Crossover number and length of SC are reduced in Ewsr1 CKO.** (A) Proportion of *Ewsr1* CKO and control spermatocytes lacking XY bivalent crossover. ** *p* < 0.01 by Student’s *t*-test. (B) Proportion of CKO and control spermatocytes lacking at least one autosomal bivalent crossover. *** *p* < 0.001 by Student’s *t*-test. (C) Upper panels, MLH1 (green) and SYCP3 (magenta) staining (same as Fig. 5A, upper panel), Lower panels, FISH for chromosome 1 (green) and chromosome 11 (magenta). Scale bar, 10 µm. (D) Proportion of spermatocytes with different numbers of crossovers (0-3) on chromosomes 1 and 11 in control and CKO. *** *p* < 0.001 by *Chi*-square test. (E) Distribution of the relative intercrossover distances on chromosome 1 and 11 measured as a fraction of the chromosomal length. *p* = 0.26 on chromosome 1, *p* = 0.66 on chromosome 11 by Student’s *t*-test. (F) Cumulative fraction plots of crossovers in the spermatocytes with double crossovers (upper panels) and single crossovers (lower panels) on chromosome 1 and chromosome 11. *Ewsr1* control, red dots; *Ewsr1* CKO, blue dots. Bars in (A)-(E) represent mean ± SD.

## References

1. Kleckner N. Meiosis: How could it work? Proceedings of the National Academy of Sciences of the United States of America. 1996;93(16):8167–74. doi: DOI 10.1073/pnas.93.16.8167. PubMed PMID: WOS:A1996VB32500004.

2. Hassold T, Hunt P. To err (meiotically) is human: the genesis of human aneuploidy. Nature reviews Genetics. 2001;2(4):280–91. doi: 10.1038/35066065. PubMed PMID: 11283700.

3. Keeney S, Giroux CN, Kleckner N. Meiosis-specific DNA double-strand breaks are catalyzed by Spo11, a member of a widely conserved protein family. Cell. 1997;88(3):375–84. PubMed PMID: 9039264.

4. Parvanov ED, Petkov PM, Paigen K. Prdm9 controls activation of mammalian recombination hotspots. Science. 2010;327(5967):835. Epub 2010/01/02. doi: 10.1126/science.1181495. PubMed PMID: 20044538; PubMed Central PMCID: PMC2821451.

5. Baudat F, Buard J, Grey C, Fledel-Alon A, Ober C, Przeworski M, et al. PRDM9 is a major determinant of meiotic recombination hotspots in humans and mice. Science. 2010;327(5967):836–40. PubMed PMID: 20044539.

6. Myers S, Bowden R, Tumian A, Bontrop RE, Freeman C, MacFie TS, et al. Drive against hotspot motifs in primates implicates the PRDM9 gene in meiotic recombination. Science. 2010;327(5967):876–9. PubMed PMID: 20044541.

7. Powers NR, Parvanov ED, Baker CL, Walker M, Petkov PM, Paigen K. The Meiotic Recombination Activator PRDM9 Trimethylates Both H3K36 and H3K4 at Recombination Hotspots In Vivo. PLoS Genet. 2016;12(6):e1006146. doi: 10.1371/journal.pgen.1006146. PubMed PMID: 27362481; PubMed Central PMCID: PMC4928815.

8. Berg IL, Neumann R, Lam KW, Sarbajna S, Odenthal-Hesse L, May CA, et al. PRDM9 variation strongly influences recombination hot-spot activity and meiotic instability in humans. Nat Genet. 2010;42(10):859–63. PubMed PMID: 20818382.

9. Romanienko PJ, Camerini-Otero RD. Cloning, characterization, and localization of mouse and human SPO11. Genomics. 1999;61(2):156–69. doi: 10.1006/geno.1999.5955. PubMed PMID: 10534401.

10. Hunter N, Kleckner N. The single-end invasion: an asymmetric intermediate at the double-strand break to double-holliday junction transition of meiotic recombination. Cell. 2001;106(1):59–70. PubMed PMID: 11461702.

11. Borner GV, Kleckner N, Hunter N. Crossover/noncrossover differentiation, synaptonemal complex formation, and regulatory surveillance at the leptotene/zygotene transition of meiosis. Cell. 2004;117(1):29–45. PubMed PMID: 15066280.

12. Martini E, Borde V, Legendre M, Audic S, Regnault B, Soubigou G, et al. Genome-wide analysis of heteroduplex DNA in mismatch repair-deficient yeast cells reveals novel properties of meiotic recombination pathways. PLoS genetics. 2011;7(9):e1002305. doi: 10.1371/journal.pgen.1002305. PubMed PMID: 21980306; PubMed Central PMCID: PMC3183076.

13. Tang S, Wu MK, Zhang R, Hunter N. Pervasive and essential roles of the Top3-Rmi1 decatenase orchestrate recombination and facilitate chromosome segregation in meiosis. Mol Cell. 2015;57(4):607–21. doi: 10.1016/j.molcel.2015.01.021. PubMed PMID: 25699709; PubMed Central PMCID: PMC4791043.

14. Baker CL, Walker M, Kajita S, Petkov PM, Paigen K. PRDM9 binding organizes hotspot nucleosomes and limits Holliday junction migration. Genome research. 2014;24(5):724–32. doi: 10.1101/gr.170167.113. PubMed PMID: 24604780; PubMed Central PMCID: PMC4009602.

15. Kauppi L, Barchi M, Lange J, Baudat F, Jasin M, Keeney S. Numerical constraints and feedback control of double-strand breaks in mouse meiosis. Genes & development. 2013;27(8):873–86. doi: 10.1101/gad.213652.113. PubMed PMID: 23599345; PubMed Central PMCID: PMC3650225.

16. Cole F, Kauppi L, Lange J, Roig I, Wang R, Keeney S, et al. Homeostatic control of recombination is implemented progressively in mouse meiosis. Nat Cell Biol. 2012;14(4):424–30. doi: 10.1038/ncb2451. PubMed PMID: 22388890; PubMed Central PMCID: PMC3319518.

17. Koehler KE, Cherry JP, Lynn A, Hunt PA, Hassold TJ. Genetic control of mammalian meiotic recombination. I. Variation in exchange frequencies among males from inbred mouse strains. Genetics. 2002;162(1):297–306. PubMed PMID: 12242241; PubMed Central PMCID: PMC1462263.

18. Baker CL, Kajita S, Walker M, Saxl RL, Raghupathy N, Choi K, et al. PRDM9 drives evolutionary erosion of hotspots in Mus musculus through haplotype-specific initiation of meiotic recombination. PLoS genetics. 2015;11(1):e1004916. doi: 10.1371/journal.pgen.1004916. PubMed PMID: 25568937; PubMed Central PMCID: PMC4287450.

19. Diagouraga B, Clement JAJ, Duret L, Kadlec J, de Massy B, Baudat F. PRDM9 Methyltransferase Activity Is Essential for Meiotic DNA Double-Strand Break Formation at Its Binding Sites. Molecular cell. 2018. doi: 10.1016/j.molcel.2018.01.033. PubMed PMID: 29478809.

20. Walker M, Billings T, Baker CL, Powers N, Tian H, Saxl RL, et al. Affinity-seq detects genome-wide PRDM9 binding sites and reveals the impact of prior chromatin modifications on mammalian recombination hotspot usage. Epigenetics Chromatin. 2015;8:31. doi: 10.1186/s13072-015-0024-6. PubMed PMID: 26351520; PubMed Central PMCID: PMC4562113.

21. Hunter N. Meiotic Recombination: The Essence of Heredity. Cold Spring Harbor perspectives in biology. 2015;7(12). doi: 10.1101/cshperspect.a016618. PubMed PMID: 26511629.

22. Robert T, Nore A, Brun C, Maffre C, Crimi B, Bourbon HM, et al. The TopoVIB-Like protein family is required for meiotic DNA double-strand break formation. Science. 2016;351(6276):943–9. doi: 10.1126/science.aad5309. PubMed PMID: 26917764.

23. Kumar R, Ghyselinck N, Ishiguro K, Watanabe Y, Kouznetsova A, Hoog C, et al. MEI4 - a central player in the regulation of meiotic DNA double-strand break formation in the mouse. J Cell Sci. 2015;128(9):1800–11. Epub 2015/03/22. doi: 10.1242/jcs.165464. PubMed PMID: 25795304.

24. Stanzione M, Baumann M, Papanikos F, Dereli I, Lange J, Ramlal A, et al. Meiotic DNA break formation requires the unsynapsed chromosome axis-binding protein IHO1 (CCDC36) in mice. Nat Cell Biol. 2016;18(11):1208–20. Epub 2016/10/28. doi: 10.1038/ncb3417. PubMed PMID: 27723721; PubMed Central PMCID: PMCPMC5089853.

25. Llano E, Herran Y, Garcia-Tunon I, Gutierrez-Caballero C, de Alava E, Barbero JL, et al. Meiotic cohesin complexes are essential for the formation of the axial element in mice. J Cell Biol. 2012;197(7):877–85. Epub 2012/06/20. doi: 10.1083/jcb.201201100. PubMed PMID: 22711701; PubMed Central PMCID: PMC3384418.

26. Ishiguro K, Kim J, Fujiyama-Nakamura S, Kato S, Watanabe Y. A new meiosis-specific cohesin complex implicated in the cohesin code for homologous pairing. EMBO reports. 2011;12(3):267–75. doi: 10.1038/embor.2011.2. PubMed PMID: 21274006; PubMed Central PMCID: PMC3059921.

27. Parvanov ED, Tian H, Billings T, Saxl RL, Spruce C, Aithal R, et al. PRDM9 interactions with other proteins provide a link between recombination hotspots and the chromosomal axis in meiosis. Mol Biol Cell. 2017;28(3):488–99. doi: 10.1091/mbc.E16-09-0686. PubMed PMID: 27932493; PubMed Central PMCID: PMC5341731.

28. Li H, Watford W, Li C, Parmelee A, Bryant MA, Deng C, et al. Ewing sarcoma gene EWS is essential for meiosis and B lymphocyte development. The Journal of clinical investigation. 2007;117(5):1314–23. doi: 10.1172/JCI31222. PubMed PMID: 17415412; PubMed Central PMCID: PMC1838927.

29. Hayashi K, Yoshida K, Matsui Y. A histone H3 methyltransferase controls epigenetic events required for meiotic prophase. Nature. 2005;438(7066):374-8. PubMed PMID: 16292313.

30. Cobb J, Cargile B, Handel MA. Acquisition of competence to condense metaphase I chromosomes during spermatogenesis. Developmental biology. 1999;205(1):49–64. doi: 10.1006/dbio.1998.9101. PubMed PMID: 9882497.

31. Mahadevaiah SK, Turner JM, Baudat F, Rogakou EP, de Boer P, Blanco-Rodriguez J, et al. Recombinational DNA double-strand breaks in mice precede synapsis. Nature genetics. 2001;27(3):271–6. doi: 10.1038/85830. PubMed PMID: 11242108.

32. Sun F, Fujiwara Y, Reinholdt LG, Hu J, Saxl RL, Baker CL, et al. Nuclear localization of PRDM9 and its role in meiotic chromatin modifications and homologous synapsis. Chromosoma. 2015. doi: 10.1007/s00412-015-0511-3. PubMed PMID: 25894966.

33. Buard J, Barthes P, Grey C, de Massy B. Distinct histone modifications define initiation and repair of meiotic recombination in the mouse. Embo J. 2009;28(17):2616–24. PubMed PMID: 19644444.

34. Altemose N, Noor N, Bitoun E, Tumian A, Imbeault M, Chapman JR, et al. A map of human PRDM9 binding provides evidence for novel behaviors of PRDM9 and other zinc-finger proteins in meiosis. eLife. 2017;6. doi: 10.7554/eLife.28383. PubMed PMID: 29072575.

35. Baker CL, Petkova P, Walker M, Flachs P, Mihola O, Trachtulec Z, et al. Multimer Formation Explains Allelic Suppression of PRDM9 Recombination Hotspots. PLoS genetics. 2015;11(9):e1005512. doi: 10.1371/journal.pgen.1005512. PubMed PMID: 26368021; PubMed Central PMCID: PMC4569383.

36. Patel A, Horton JR, Wilson GG, Zhang X, Cheng X. Structural basis for human PRDM9 action at recombination hot spots. Genes & development. 2016;30(3):257–65. doi: 10.1101/gad.274928.115. PubMed PMID: 26833727; PubMed Central PMCID: PMC4743056.

37. Billings T, Parvanov ED, Baker CL, Walker M, Paigen K, Petkov PM. DNA binding specificities of the long zinc-finger recombination protein PRDM9. Genome Biol. 2013;14(4):R35. doi: 10.1186/gb-2013-14-4-r35. PubMed PMID: 23618393; PubMed Central PMCID: PMC4053984.

38. Cohen PE, Pollack SE, Pollard JW. Genetic analysis of chromosome pairing, recombination, and cell cycle control during first meiotic prophase in mammals. Endocrine reviews. 2006;27(4):398–426. doi: 10.1210/er.2005-0017. PubMed PMID: 16543383.

39. Brick K, Smagulova F, Khil P, Camerini-Otero RD, Petukhova GV. Genetic recombination is directed away from functional genomic elements in mice. Nature. 2012;485(7400):642-5. Epub 2012/06/05. doi: 10.1038/nature11089. PubMed PMID: 22660327; PubMed Central PMCID: PMC3367396.

40. Smagulova F, Gregoretti IV, Brick K, Khil P, Camerini-Otero RD, Petukhova GV. Genome-wide analysis reveals novel molecular features of mouse recombination hotspots. Nature. 2011;472(7343):375-8. Epub 2011/04/05. doi: 10.1038/nature09869. PubMed PMID: 21460839; PubMed Central PMCID: PMCPMC3117304.

41. Lange J, Yamada S, Tischfield SE, Pan J, Kim S, Zhu X, et al. The Landscape of Mouse Meiotic Double-Strand Break Formation, Processing, and Repair. Cell. 2016;167(3):695–708 e16. Epub 2016/10/22. doi: 10.1016/j.cell.2016.09.035. PubMed PMID: 27745971; PubMed Central PMCID: PMCPMC5117687.

42. Anderson LK, Reeves A, Webb LM, Ashley T. Distribution of crossing over on mouse synaptonemal complexes using immunofluorescent localization of MLH1 protein. Genetics. 1999;151(4):1569–79. PubMed PMID: 10101178.

43. Baier B, Hunt P, Broman KW, Hassold T. Variation in Genome-Wide Levels of Meiotic Recombination Is Established at the Onset of Prophase in Mammalian Males. PLoS genetics. 2014;10(1). doi: ARTN e1004125 10.1371/journal.pgen.1004125. PubMed PMID: WOS:000336525000068.

44. Gruhn JR, Rubio C, Broman KW, Hunt PA, Hassold T. Cytological studies of human meiosis: sex-specific differences in recombination originate at, or prior to, establishment of double-strand breaks. PloS one. 2013;8(12):e85075. doi: 10.1371/journal.pone.0085075. PubMed PMID: 24376867; PubMed Central PMCID: PMC3869931.

45. Wang S, Hassold T, Hunt P, White MA, Zickler D, Kleckner N, et al. Inefficient Crossover Maturation Underlies Elevated Aneuploidy in Human Female Meiosis. Cell. 2017. doi: 10.1016/j.cell.2017.02.002. PubMed PMID: 28262352.

46. Lynn A, Koehler KE, Judis L, Chan ER, Cherry JP, Schwartz S, et al. Covariation of synaptonemal complex length and mammalian meiotic exchange rates. Science. 2002;296(5576):2222-5. PubMed PMID: 12052900.

47. Petkov PM, Broman KW, Szatkiewicz JP, Paigen K. Crossover interference underlies sex differences in recombination rates. Trends Genet. 2007;23(11):539–42. Epub 2007/10/30. doi: 10.1016/j.tig.2007.08.015. PubMed PMID: 17964681.

48. de Boer E, Stam P, Dietrich AJ, Pastink A, Heyting C. Two levels of interference in mouse meiotic recombination. Proc Natl Acad Sci U S A. 2006;103(25):9607–12. doi: 10.1073/pnas.0600418103. PubMed PMID: 16766662; PubMed Central PMCID: PMC1475796.

49. Imai Y, Baudat F, Taillepierre M, Stanzione M, Toth A, de Massy B. The PRDM9 KRAB domain is required for meiosis and involved in protein interactions. Chromosoma. 2017;126(6):681–95. Epub 2017/05/21. doi: 10.1007/s00412-017-0631-z. PubMed PMID: 28527011; PubMed Central PMCID: PMCPMC5688218.

50. Tian H, Billings T, Petkov PM. CXXC1 is redundant for normal DNA double-strand break formation and meiotic recombination in mouse. BIORXIV preprint server. 2018. doi: doi: https://doi.org/10.1101/305508.

51. Schwartz JC, Cech TR, Parker RR. Biochemical Properties and Biological Functions of FET Proteins. Annual review of biochemistry. 2015;84:355–79. doi: 10.1146/annurev-biochem-060614-034325. PubMed PMID: 25494299.

52. Fisher C. The diversity of soft tissue tumours with EWSR1 gene rearrangements: a review. Histopathology. 2014;64(1):134–50. Epub 2013/12/11. doi: 10.1111/his.12269. PubMed PMID: 24320889.

53. Imai Y, Baudat F, Taillepierre M, Stanzione M, Toth A, de Massy B. The PRDM9 KRAB domain is required for meiosis and involved in protein interactions. Chromosoma. 2017. doi: 10.1007/s00412-017-0631-z. PubMed PMID: 28527011.

54. Mulligan P, Westbrook TF, Ottinger M, Pavlova N, Chang B, Macia E, et al. CDYL bridges REST and histone methyltransferases for gene repression and suppression of cellular transformation. Mol Cell. 2008;32(5):718–26. doi: 10.1016/j.molcel.2008.10.025. PubMed PMID: 19061646.

55. Kohler S, Wojcik M, Xu K, Dernburg AF. Superresolution microscopy reveals the three-dimensional organization of meiotic chromosome axes in intact Caenorhabditis elegans tissue. Proceedings of the National Academy of Sciences of the United States of America. 2017;114(24):E4734-E43. doi: 10.1073/pnas.1702312114. PubMed PMID: 28559338; PubMed Central PMCID: PMC5474826.

56. Kugou K, Fukuda T, Yamada S, Ito M, Sasanuma H, Mori S, et al. Rec8 guides canonical Spo11 distribution along yeast meiotic chromosomes. Molecular biology of the cell. 2009;20(13):3064–76. doi: 10.1091/mbc.E08-12-1223. PubMed PMID: 19439448; PubMed Central PMCID: PMC2704158.

57. Ward A, Hopkins J, McKay M, Murray S, Jordan PW. Genetic Interactions Between the Meiosis-Specific Cohesin Components, STA G3, REC8, and RAD21L. G3. 2016;6(6):1713-24. doi: 10.1534/g3.116.029462. PubMed PMID: 27172213; PubMed Central PMCID: PMC4889667.

58. Bannister LA, Reinholdt LG, Munroe RJ, Schimenti JC. Positional cloning and characterization of mouse mei8, a disrupted allelle of the meiotic cohesin Rec8. Genesis. 2004;40(3):184–94. doi: 10.1002/gene.20085. PubMed PMID: 15515002.

59. Xu H, Beasley MD, Warren WD, van der Horst GT, McKay MJ. Absence of mouse REC8 cohesin promotes synapsis of sister chromatids in meiosis. Developmental cell. 2005;8(6):949–61. doi: 10.1016/j.devcel.2005.03.018. PubMed PMID: 15935783.

60. Brar GA, Kiburz BM, Zhang Y, Kim JE, White F, Amon A. Rec8 phosphorylation and recombination promote the step-wise loss of cohesins in meiosis. Nature. 2006;441(7092):532-6. doi: 10.1038/nature04794. PubMed PMID: 16672979.

61. Buonomo SB, Clyne RK, Fuchs J, Loidl J, Uhlmann F, Nasmyth K. Disjunction of homologous chromosomes in meiosis I depends on proteolytic cleavage of the meiotic cohesin Rec8 by separin. Cell. 2000;103(3):387–98. PubMed PMID: 11081626.

62. Eijpe M, Offenberg H, Jessberger R, Revenkova E, Heyting C. Meiotic cohesin REC8 marks the axial elements of rat synaptonemal complexes before cohesins SMC1beta and SMC3. The Journal of cell biology. 2003;160(5):657–70. doi: 10.1083/jcb.200212080. PubMed PMID: 12615909; PubMed Central PMCID: PMC2173354.

63. Lee J, Iwai T, Yokota T, Yamashita M. Temporally and spatially selective loss of Rec8 protein from meiotic chromosomes during mammalian meiosis. Journal of cell science. 2003;116(Pt 13):2781–90. doi: 10.1242/jcs.00495. PubMed PMID: 12759374.

64. Yoon SW, Lee MS, Xaver M, Zhang L, Hong SG, Kong YJ, et al. Meiotic prophase roles of Rec8 in crossover recombination and chromosome structure. Nucleic acids research. 2016;44(19):9296–314. doi: 10.1093/nar/gkw682. PubMed PMID: 27484478; PubMed Central PMCID: PMC5100558.

65. Reynolds A, Qiao HY, Yang Y, Chen JK, Jackson N, Biswas K, et al. RNF212 is a dosage-sensitive regulator of crossing-over during mammalian meiosis. Nature genetics. 2013;45(3):269–78. doi: 10.1038/ng.2541. PubMed PMID: WOS:000315664800010.

66. Holloway JK, Sun XF, Yokoo R, Villeneuve AM, Cohen PE. Mammalian CNTD1 is critical for meiotic crossover maturation and deselection of excess precrossover sites. Journal of Cell Biology. 2014;205(5):633–41. doi: 10.1083/jcb.201401122. PubMed PMID: WOS:000337210800005.

67. Kneitz B, Cohen PE, Avdievich E, Zhu LY, Kane MF, Hou H, et al. MutS homolog 4 localization to meiotic chromosomes is required for chromosome pairing during meiosis in male and female mice. Genes & development. 2000;14(9):1085–97. PubMed PMID: WOS:000087101500006.

68. Berchowitz LE, Copenhaver GP. Genetic Interference: Don’t Stand So Close to Me. Curr Genomics. 2010;11(2):91–102. doi: Doi 10.2174/138920210790886835. PubMed PMID: WOS:000275150100002.

69. Paigen K, Szatkiewicz JP, Sawyer K, Leahy N, Parvanov ED, Ng SH, et al. The recombinational anatomy of a mouse chromosome. PLoS genetics. 2008;4(7):e1000119. doi: 10.1371/journal.pgen.1000119. PubMed PMID: 18617997; PubMed Central PMCID: PMC2440539.

70. Billings T, Sargent EE, Szatkiewicz JP, Leahy N, Kwak IY, Bektassova N, et al. Patterns of recombination activity on mouse chromosome 11 revealed by high resolution mapping. PLoS One. 2010;5(12):e15340. doi: 10.1371/journal.pone.0015340. PubMed PMID: 21170346; PubMed Central PMCID: PMC2999565.

71. Liu EY, Morgan AP, Chesler EJ, Wang W, Churchill GA, Pardo-Manuel de Villena F. High-resolution sex-specific linkage maps of the mouse reveal polarized distribution of crossovers in male germline. Genetics. 2014;197(1):91–106. Epub 2014/03/01. doi: 10.1534/genetics.114.161653. PubMed PMID: 24578350; PubMed Central PMCID: PMC4012503.

72. Carofiglio F, Inagaki A, de Vries S, Wassenaar E, Schoenmakers S, Vermeulen C, et al. SPO11-independent DNA repair foci and their role in meiotic silencing. PLoS genetics. 2013;9(6):e1003538. doi: 10.1371/journal.pgen.1003538. PubMed PMID: 23754961; PubMed Central PMCID: PMC3675022.

73. Agostinho A, Manneberg O, van Schendel R, Hernandez-Hernandez A, Kouznetsova A, Blom H, et al. High density of REC8 constrains sister chromatid axes and prevents illegitimate synaptonemal complex formation. EMBO reports. 2016;17(6):901–13. doi: 10.15252/embr.201642030. PubMed PMID: 27170622; PubMed Central PMCID: PMC5278604.

74. Bhattacharyya T, Walker M, Powers NR, Brunton C, Fine AD, Petkov PM, et al. Prdm9 and meiotic cohesin proteins 1 cooperatively promote DNA double-strand break formation in mammalian spermatocytes. Current biology. 2019;29:1–17. doi: https://doi.org/10.1016/j.cub.2019.02.007. PubMed Central PMCID: PMCPMC6544150.

75. Bradley A, Anastassiadis K, Ayadi A, Battey JF, Bell C, Birling MC, et al. The mammalian gene function resource: the International Knockout Mouse Consortium. Mammalian genome: official journal of the International Mammalian Genome Society. 2012;23(9-10):580–6. doi: 10.1007/s00335-012-9422-2. PubMed PMID: 22968824; PubMed Central PMCID: PMC3463800.

76. Peters AH, Plug AW, van Vugt MJ, de Boer P. A drying-down technique for the spreading of mammalian meiocytes from the male and female germline. Chromosome research: an international journal on the molecular, supramolecular and evolutionary aspects of chromosome biology. 1997;5(1):66–8. PubMed PMID: 9088645.

77. Davies B, Hatton E, Altemose N, Hussin JG, Pratto F, Zhang G, et al. Re-engineering the zinc fingers of PRDM9 reverses hybrid sterility in mice. Nature. 2016;530(7589):171-6. doi: 10.1038/nature16931. PubMed PMID: 26840484; PubMed Central PMCID: PMC4756437.

78. Jee J, Rozowsky J, Yip KY, Lochovsky L, Bjornson R, Zhong G, et al. ACT: aggregation and correlation toolbox for analyses of genome tracks. Bioinformatics. 2011;27(8):1152–4. doi: 10.1093/bioinformatics/btr092. PubMed PMID: 21349863; PubMed Central PMCID: PMC3072554.

79. Khil PP, Smagulova F, Brick KM, Camerini-Otero RD, Petukhova GV. Sensitive mapping of recombination hotspots using sequencing-based detection of ssDNA. Genome research. 2012;22(5):957–65. doi: 10.1101/gr.130583.111. PubMed PMID: 22367190; PubMed Central PMCID: PMC3337440.

