## Supplementary figures and images for "EWSR1 affects PRDM9-dependent histone 3 methylation and provides a link between recombination hotspots and the chromosome axis"

### Supplemental figure 1

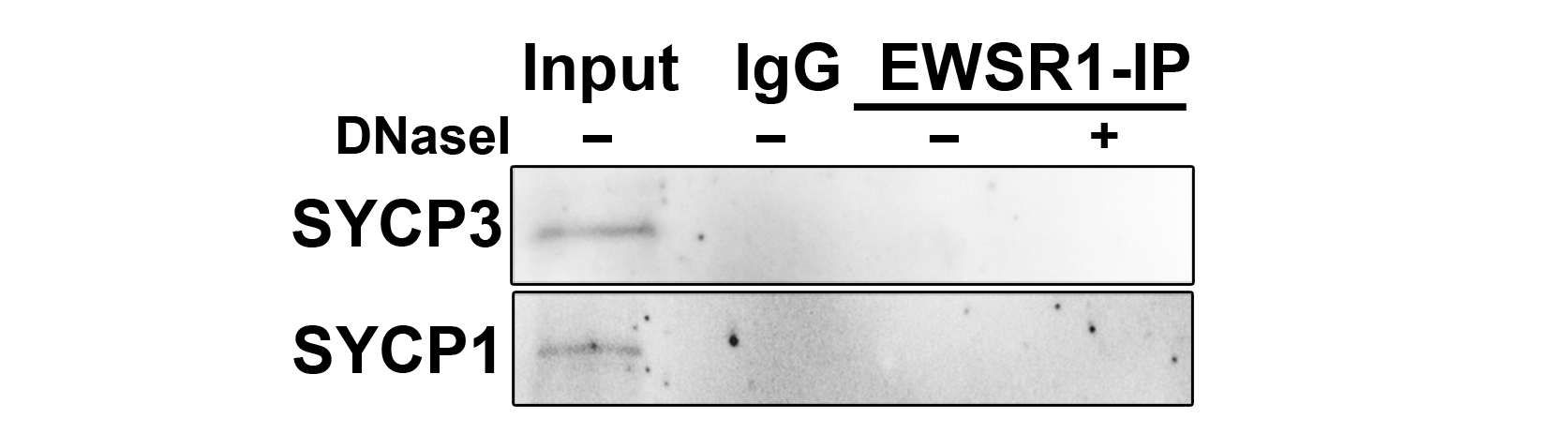

### Supplemental figure 2

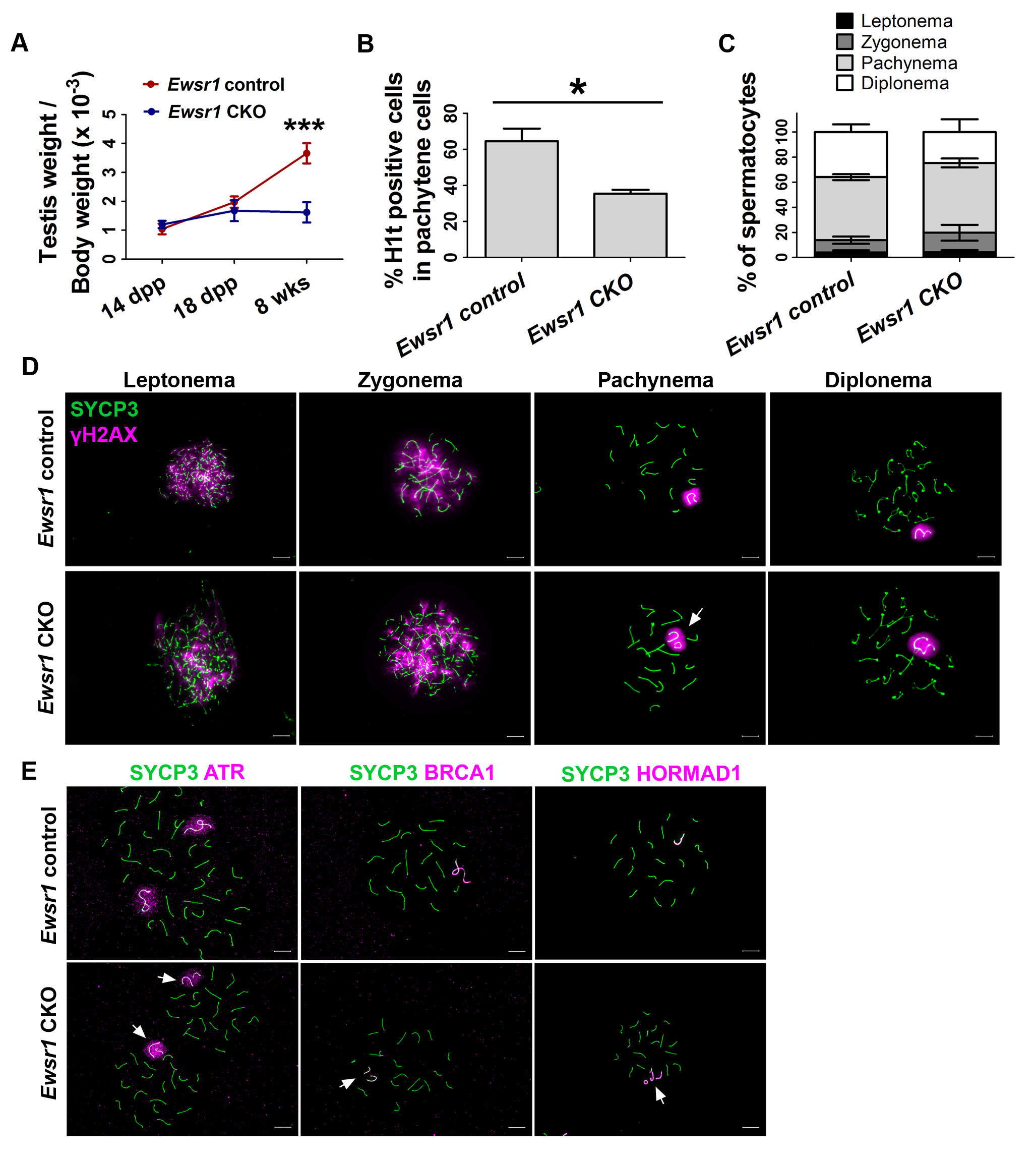

### Supplemental figure 3

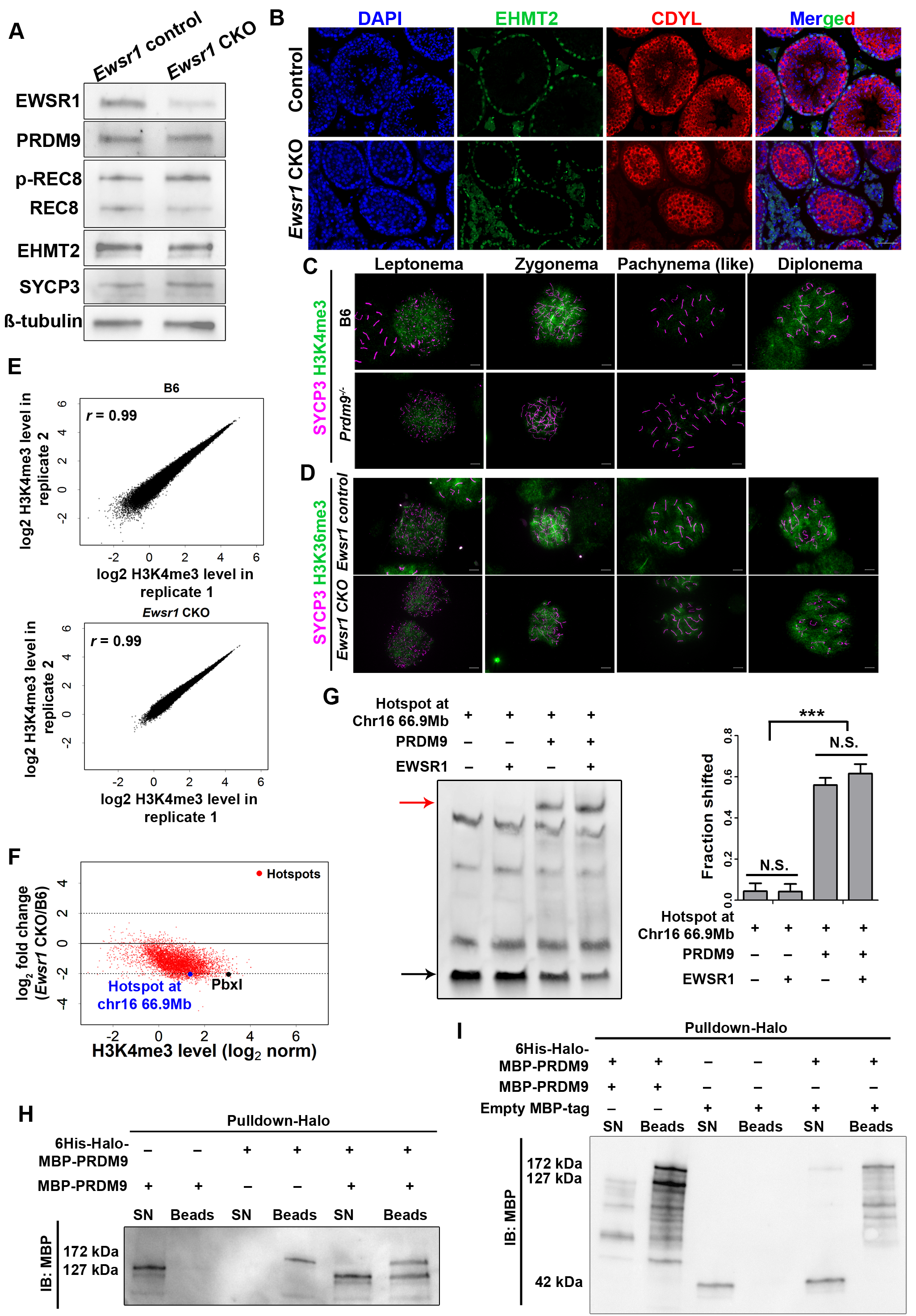

### Supplemental figure 4

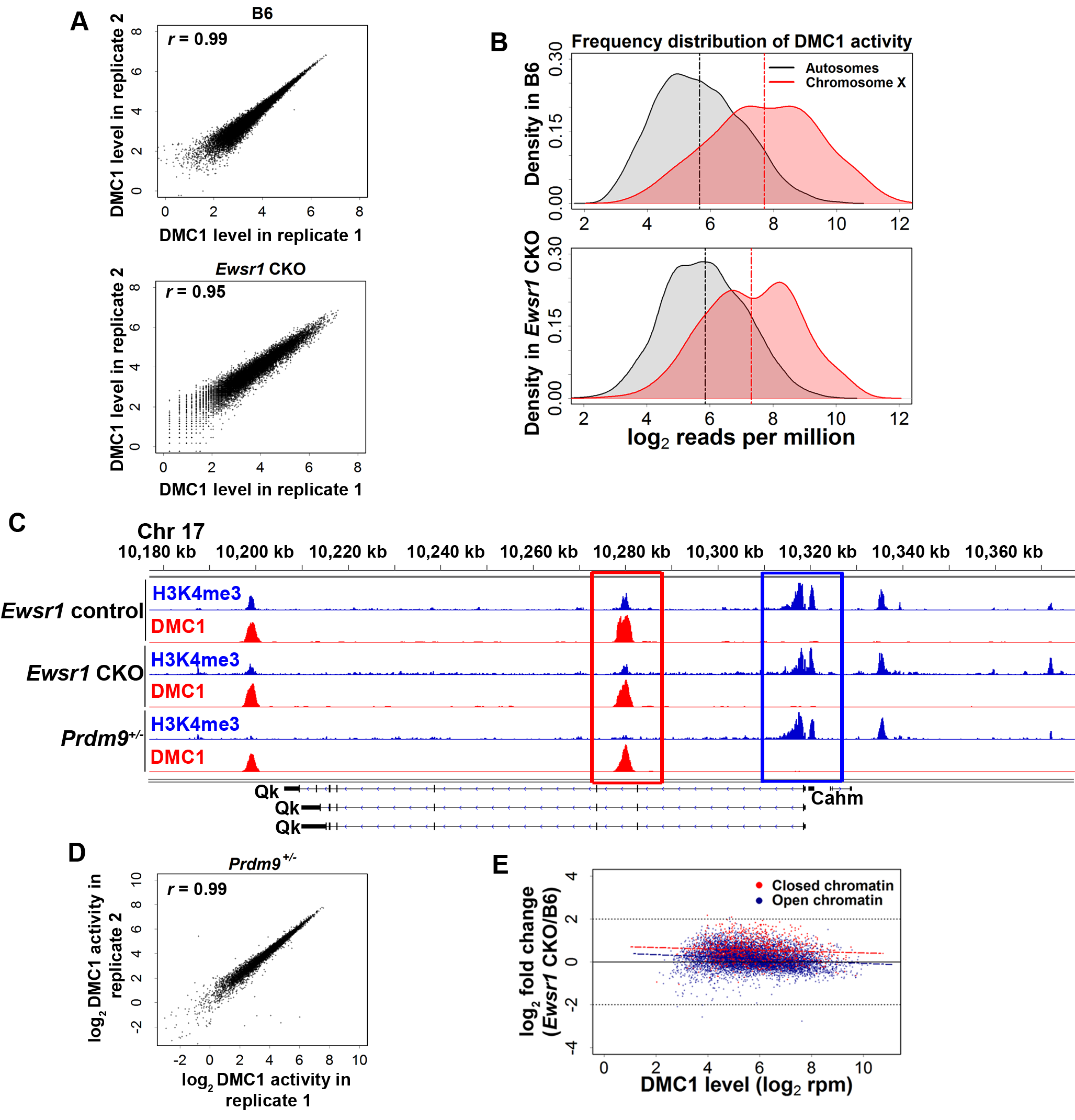

### Supplemental figure 5

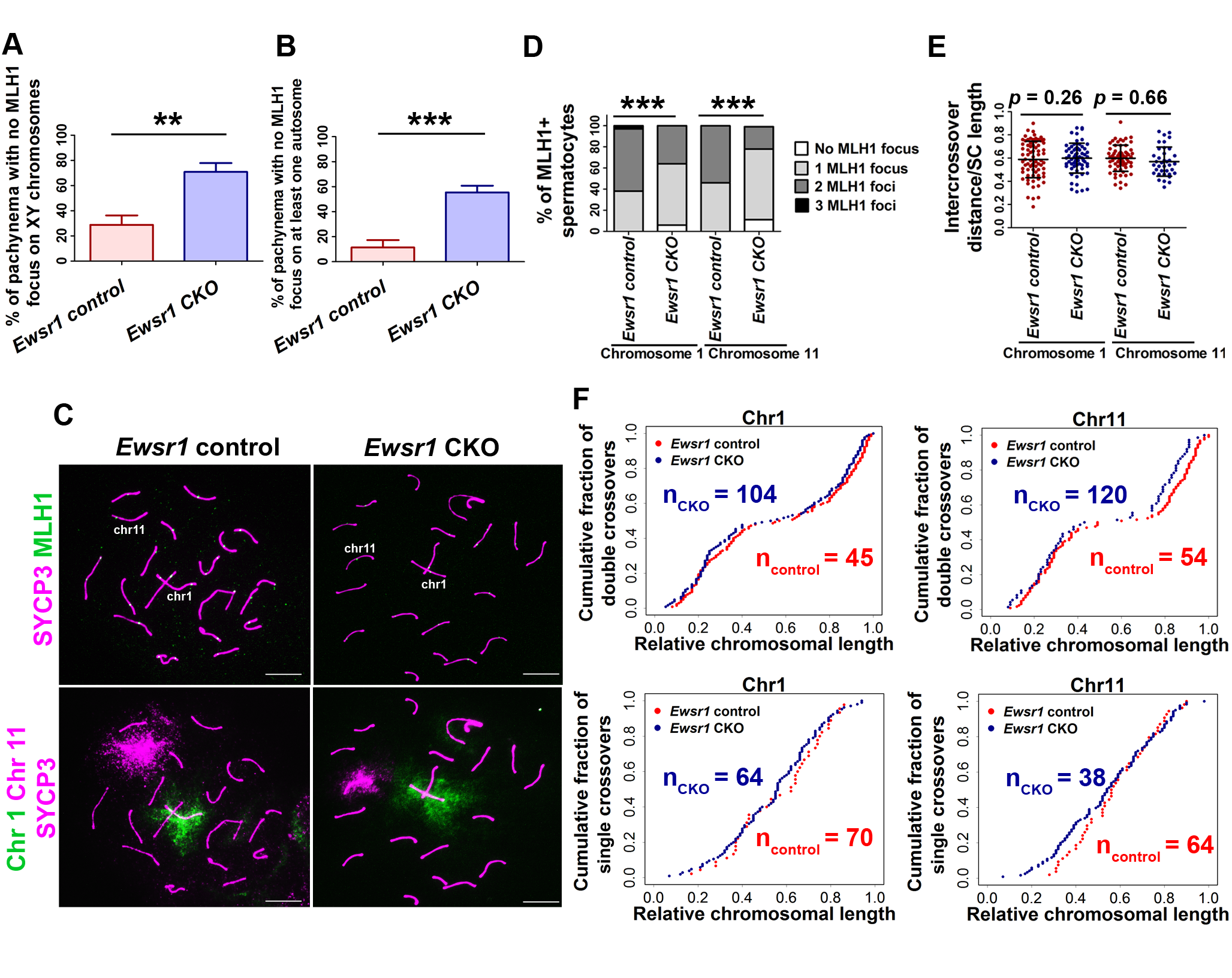
